# In toto analysis of multicellular arrangement reduces embryonic tissue diversity to two archetypes that require specific cadherin expression

**DOI:** 10.1101/2024.12.06.626808

**Authors:** Max Brambach, Marvin Albert, Jérôme Julmi, Robert Bill, Darren Gilmour

## Abstract

Breakthroughs in transcriptomics are providing exciting new ways to characterise cellular diversity within organisms^1–3^. Yet, organisms are more than the sum of their differentiated cells, as the biological function of most cell types only emerges when they are organised into tissues with characteristic architectures^4^. The mechanisms that drive architectural diversification at the tissue level remain poorly understood, in part due to a general lack of methods for directly comparing different organisational patterns. Here we establish nuQLOUD, an efficient imaging and computational framework that reduces complex tissues to clouds of nuclear positions, enabling the extraction of cell-type agnostic architectural features. Applying nuQLOUD to whole developing zebrafish embryos reveals that global tissue diversity can be efficiently reduced to two primary archetypes, termed ‘amorphous’ and ‘crystalline’. We investigate the molecular drivers of tissue archetypes by focussing on cadherin cell adhesion molecules and demonstrate that the expression domains of major cadherins segregate along tissue-archetypal lines. Further spatiotemporal analysis and targeted perturbation of cadherin expression in different organs identifies N-Cadherin as a general driver of the amorphous archetype. Thus, this systematic investigation of architectural diversity provides a new way to conceptualise embryonic organisation and understand drivers of tissue diversification within a standardised framework.

## Introduction

Cells are the basic units of life. Advances in transcriptomics now enable cellular diversity within organisms to be characterised at an unprecedented resolution^1–3^. The power of this approach is that it exploits the standard language of mRNA sequences to define and compare diverse cells in a systematic and quantitative manner. When combined with perturbation experiments, such expression profiling methods can identify drivers of cell type diversification^5,6^. However, characterising cell compositions via current -omics methods is only part of the equation, as organisms are much more than the sum of their cell types. Indeed, the biological functions of most cell types only emerge when they are organised into multicellular tissues with specific architectures^4^. The barrier function of the skin depends on keratinocytes being organised in sheets, the contractile function of muscles depends on their being bundled into longitudinal fibres, the computational function of the brain depends on neurons being connected in networks, and so on. Tissues can therefore be considered the functional units of life, in metazoans at least, and these functions require proper organisation. It is therefore surprising that, while the study of tissue biology dates to Aristotle, there is still no standard language for describing, categorising and comparing tissue organisation.

An important step towards characterising architectural diversity is the development of advanced imaging methods such as light-sheet microscopy, which allow standardised imaging of whole developing organisms at single cell resolution^7,8^. Such *in toto* imaging has been combined with large-scale tracking of cell nuclei to provide insights into dynamic germ layer interactions during zebrafish gastrulation^9^ and the origins of cell fates from gastrulation to organogenesis stages of mammalian development^10^. However, while *in toto* microscopy enabled unprecedented imaging of entire animals, this breakthrough has not yet delivered a greater understanding of the architectural diversity that exists within organisms. A key limitation has been a lack of standardised frameworks for quantifying and comparing multicellular organisation. Bioimage analysis has generally focused on single cell segmentation methods that prove to be very powerful for the description of individual cell morphologies^11,12^. However, it can be challenging to extend these to complex tissues and whole organisms due to the high variability in cell morphologies and distributions^13^. This heterogeneity also underlies another challenge: descriptors that can define the shape of cells in one tissue may not be informative for another. For example, epithelial cells can often be efficiently quantified as densely packed polygons^14^, whereas ongoing efforts to define the architecture of the nervous system focus on the morphology and connectivity of axonal and dendritic protrusions^15^. Cell morphology segmentation methods encourage a focus on cell-scale features that may not be relevant to architecture at the tissue-scale, which can lead to ‘not seeing the forest for the trees’. Thus, the development of efficient frameworks that allow diverse tissue organisations to be directly compared in a ‘cell-type agnostic’ manner represents an important step towards understanding the architectural diversity that exists within and across organisms.

The lack of methods for comparing multicellular architecture has specifically hindered investigations into the genetic regulation of tissue diversification. At the cellular scale, morphogenesis is driven primarily by the actomyosin cytoskeleton and regulated by RhoA GTPases universally present in all cells. At the tissue scale, cytoskeletal activities are interconnected via cell-cell adhesion that coordinate cell shaping and generate specific multicellular structures required for diverse organ functions. This indicates that patterned expression of cell adhesion molecules, most notably cadherins, could enable genetically-regulated changes in tissue organisation. Cadherins have long been studied for their role in tissue integrity and changes in cadherin expression are hallmarks of degenerative diseases and cancer^16,17^. Differential expression of cadherins can promote tissue segregation during early development^18^ and cell sorting in ‘synthetic’ embryos^19^. Beyond segregation, patterned cadherin expression plays a key role in coordinated cell movements^20,21^ and lately a ‘cadherin code’ has been shown to increase the robustness of morphogen patterning in the central nervous system (CNS)^22^. Notably, the large expansion of the cadherin superfamily that coincided with the emergence of vertebrates - with more than 100 cadherin and cadherin-related proteins being present in humans^23^ - has been proposed to reflect the greater morphological diversification observed in these species^24^. However, it remains unclear if differences in cadherin expression can explain the different arrangements of cells that shape tissues.

In this study, we present a general approach to quantify organisational diversity by simplifying tissue complexity to the three-dimensional arrangement of constituent cells, ‘focussing on the forest rather than the trees’. By performing nuclear segmentation across entire embryos, we defined cellular positions within a common point-cloud-based framework, a standard data structure in many spatial analysis tasks, enabling comparative organisational analysis of developing tissues in a cell type-agnostic manner. This coarse-grained approach allowed us to track tissue organisation over time and correlate organisational features with gene expression. Unbiased clustering of similarly organised cells revealed that tissue heterogeneity can be reduced to two organisational ‘archetypes,’ which we termed ‘amorphous’ and ‘crystalline,’ based on criteria from material science. Defining the *in toto* expression of 12 major cadherin cell adhesion receptors showed that cadherin expression domains become bi-partitioned along tissue archetypal lines. Further investigation using spatiotemporal correlation analysis and targeted perturbations identified N-cadherin as a general regulator of the amorphous archetype.

## Results

### nuQLOUD - a framework for the embryo-wide quantification of tissue organisation based on 3D cellular arrangement

To enable the investigation of the organisational diversity of entire developing organisms, we initially focussed on zebrafish embryos at 48 hours post fertilisation (hpf), a stage where the the progenitors of several major organs are already formed^25^. Multi-view light sheet microscopy was used to generate isotropic resolution imaging data of whole embryos where nuclei were stained with DAPI and individual cells were localised using TGMM nuclear segmentation (Figure 1 a, Suppl. Fig. 1), identifying the centre of mass of individual nuclei^10^. While nuclear segmentation is routinely used to count and track cells^9,10^, here we explored its potential for quantifying tissue organisation *in toto*. We reasoned that quantifying the 3D arrangement of cells via the position of their nuclei may provide an effective and standardised method to compare conserved features of tissue architecture. Overall, we quantified the distribution of more than 4,000,000 cells in their native environment, across 34 samples, an average of 116,600 ± 9,000 cells per embryo. We next took advantage of the explicit point-cloud structure of the data to characterise the local organisation of nuclei using a three-dimensional Voronoi diagram, a spatial partitioning algorithm that assigns each point a volume of closest ‘cells’ from point distributions. Such Voronoi cells are constructed by dividing 3D space into regions consisting of all points closer to a given point than to any other point. The shape and size of such Voronoi cells consequently provide an object-based representation of local, multicellular organisation^26^. A known limitation of conventional Voronoi diagrams is that cells at the embryo’s edge can have infinite size due to a lack of neighbouring points to limit their extent, making the method unsuitable for boundary tissues like skin. To overcome this, we restricted the size of boundary Voronoi cells adaptively to their neighbourhood by placing artificial points around the perimeter of the embryo such that each local neighbourhood is smoothly extended by one cell layer (see Suppl. Fig. 2, Suppl. Note 1 for details). We used this adaptively restricted Voronoi diagram to characterise the organisational context of every nucleus in entire zebrafish embryos using fourteen engineered features, including kernel density estimations over different length scales, number of neighbours and the polarity of the Voronoi cell (Figure 1 b, Suppl. Fig. 3 a-d, Suppl. Note 2). Clustering of these features by similarity across all analysed cells identified three organisational feature classes based entirely on local nuclear position that we termed anisotropy, density, and irregularity (Figure 1 c, Suppl. Fig. 3 e,f). The distribution of these feature classes highlighted different anatomical regions in a combinatorial way and mapped to different regions of the organisational feature space (Figure 1 d, Suppl. Fig. 3 g,h), which was conserved between samples (Suppl. Fig. 4). This demonstrates that our whole-organism description of multicellular arrangement robustly captures meaningful organisational differences within one unified framework, which we term nuQLOUD (NUclear-based Quantification of Local Organisation via cellUlar Distributions).

**Figure 1:**
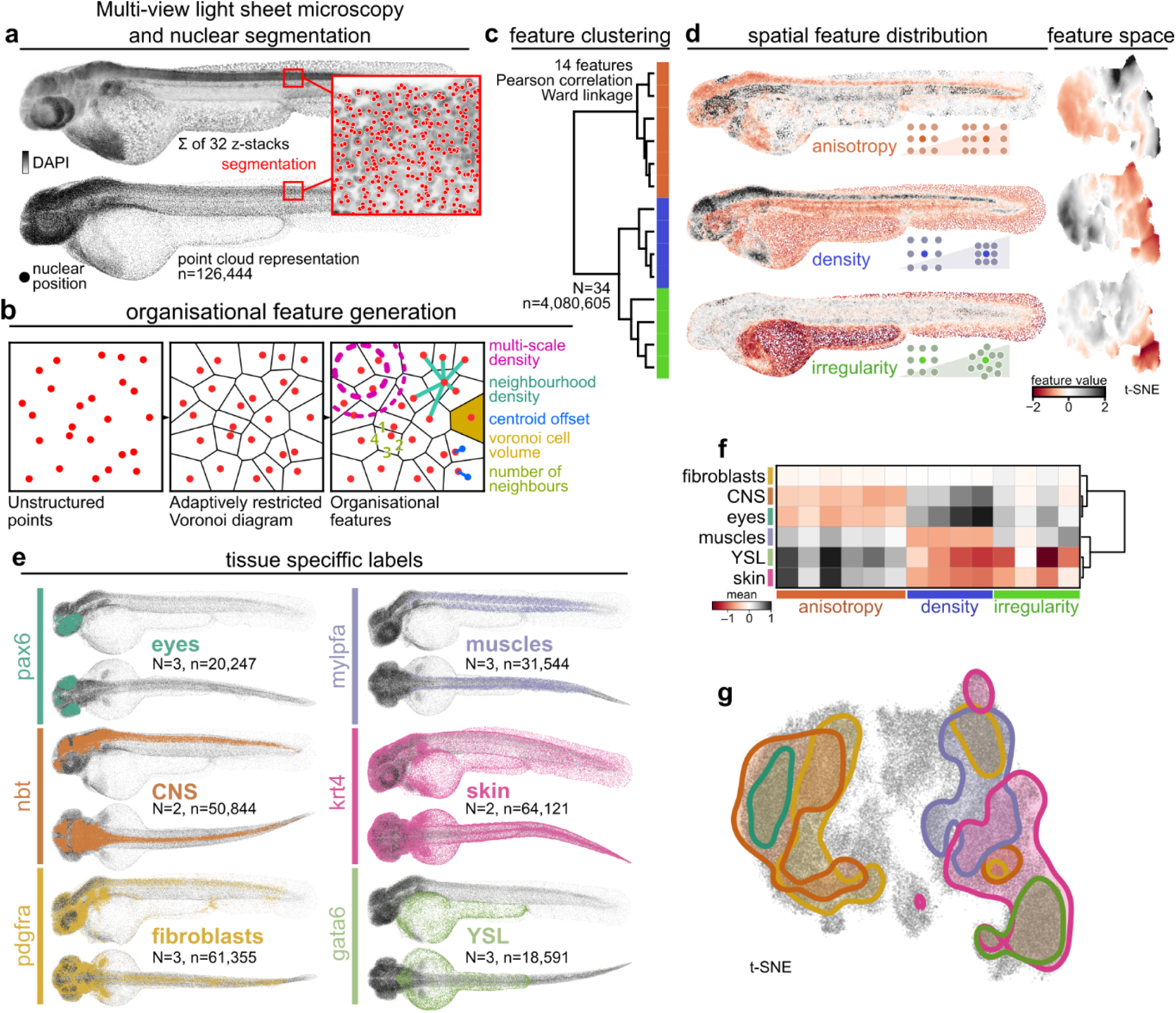
In toto quantification of multicellular organisation. a: Multi-view light sheet microscopy and nuclear segmentation transform in toto volumetric image data into point cloud representation. b: Multicellular organisation is represented for each cell by a fourteen-dimensional feature vector that describes the local point distribution. Note that explicit coordinates are not part of that feature set. c: Similarity clustering of organisational features reveals three feature classes: anisotropy, density, and irregularity. d: The spatial distributions of the organisational feature classes are distinct and highlight different anatomical regions. Moreover, feature classes exhibit different distributions on a t-SNE embedding of the organisational feature space. e: Tissue specific gene expression patterns are mapped on the point cloud representation; individual nuclei are classified as expressing/non-expressing based on fluorescence intensity of the label (HCR, transgenic line). f: Mean organisational feature profiles quantify organisational diversity. g: t-SNE embedding of organisational feature space with gene expression patterns overlayed. Feature values are z-scored across all cells of multiple samples.N: number of samples, n: total number of cells

To map out the organisational landscape of whole embryos, we used nuQLOUD to characterise and compare the organisation of major tissues, including skin, brain muscles and connective tissue (Figure 1 e, Suppl. Fig. 5 a). For that, we incorporated tissue specific gene expression information from transgenic zebrafish lines and via hybridisation chain reaction fluorescent in situ hybridisation (HCR). We classified all cells for the presence or absence of fluorescence using an adapted version of the *in toto* imaging and segmentation approach (Suppl. Fig. 6). In this way, we were able to assign organisational signatures to genetically defined tissues (Figure 1 g, Suppl. Fig. 5 b) and compare these directly (Fig. 1 f, Suppl. Fig. 5 c). This allowed projecting tissue landmarks onto the feature space representation and indicated organisational overlap between tissues that are far apart in both lineage and function, such as skin and muscles, which are derived from ectoderm and mesoderm, respectively; both exhibit low density and a high degree of anisotropy (Fig. 1 f,g, Suppl. Fig. 5 c). Interestingly, clustering of all analysed tissues based on their organisation revealed a bimodal distribution with muscles, skin and yolk syncytial layer (YSL) in one group and eyes and CNS in the other (Fig. 1 f). These results show how our nuQLOUD framework can be used to identify shared organisational patterns across vastly different organs in whole organisms.

### Unbiased classification of tissue architecture reduces global organisational diversity to two archetypes

Next, we expanded our nuQLOUD analysis from gene expression-defined organs to all cells at multiple stages of zebrafish embryogenesis to identify and analyse stereotypical organisational patterns. Cells with a shared organisational signature were grouped into eleven ‘organisational motifs’ using a Gaussian mixture model (Fig. 2 a, Suppl. Fig. 7, Suppl. Note 3). The spatial distributions of individual motifs were compact and mapped to anatomical regions when projected on individual samples, highlighting that the nuQLOUD framework was able to differentiate between tissues solely based on the local distribution of nuclei (Suppl. Fig. 8). We tracked the organisational motifs between 12 and 72 hpf and found that some motifs were present at all time points, while others were absent early and developed over time (Fig. 2 a,b). One example for a maintained motif was motif IX, which consistently mapped to the YSL from 12 hpf. However, cells of the median fin fold were classified in this motif from 48 hpf onward, highlighting an organisational similarity between these tissues (Fig. 2 b IX). Motif I evolves from being essentially absent at 12 hpf, to highlighting cells of the spinal cord and hindbrain at 24 hpf before spreading to include cells of the forebrain and eyes by 48 hpf, indicating that in different CNS regions cells adopt this organisation in a temporal sequence (Fig. 2 b I). This demonstrates that organisational complexity increases during early development and highlights that nuQLOUD is able to detect organisational patterns without gene expression information in an unbiased way.

**Figure 2:**
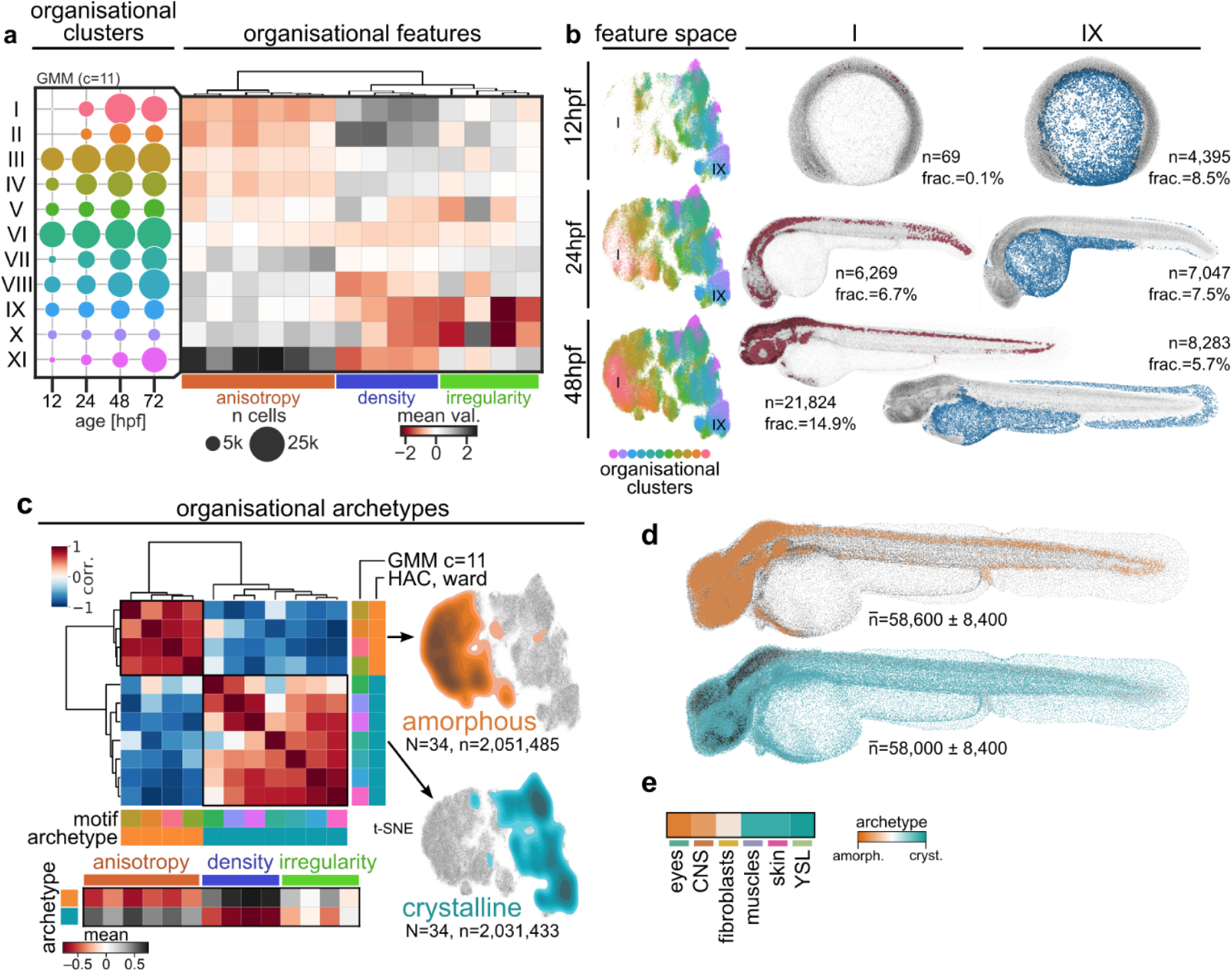
Global organisational heterogeneity can be reduced to two organisational archetypes. a,b: Organisational diversity increases over time. Organisational feature space is stratified into eleven ‘organisational motifs’ using a Gaussian Mixture Model (GMM). Using data from four different developmental time points, reveals that some motifs exist throughout early development (e.g. III, VI, IX) and others develop over time (e.g. I, II, VII) (a). Organisational features values are z-scored, and clustering is performed on data of all time points simultaneously. b illustrates two examples of organisational motifs. Motif I is absent early and cells start organising in this motif from 24 hpf on eventually covering the CNS; motif IX consistently maps to the YSL and at 48 hpf also included the median fin fold. Feature space representation (t-SNE) generated from all time points. n: number of cells per motif for shown sample. c-e: Architectural diversity can be reduced to two organisational archetypes. c Hierarchical agglomerative clustering (HAC) of organisational motifs based on tissue organisation identifies two ‘archetypes’, which map to opposing sides of organisational feature space and are defined by converse organisational features. Cells of the ‘amorphous’ archetype organise dense, isotropic, and irregular, while ‘crystalline’ cells organise low-density, anisotropic and regular. 48 hpf data was used for this clustering. Feature values are z-scored. HAC using Euclidean distance and Ward linkage. d,e: organisational archetypes at 48 hpf spatially map to cohesive regions. Cells of the CNS, sensory organs and pectoral fin buds organise amorphously, while muscles, skin and YSL are organised crystalline. Cells expressing tissue specific markers in e associate with one or the other archetype, except fibroblasts. n: mean number of cells per motif across N=34 samples ± standard deviation of the mean.

To understand higher level patterns of tissue organisation, we investigated how organisational motifs relate to each other. Clustering motifs based on their organisational similarity revealed two classes, which we term organisational ‘archetypes’, each accounting for approximately 50% of cells in the embryo at 48 hpf. Based on their organisational profile, we found that the two archetypes fit with the definitions of ‘crystalline’ and ‘amorphous’ from material sciences (Fig. 2 c). Tissues assigned to the amorphous archetype showed a dense, isotropic and irregular arrangement of nuclei and included cells of the eyes, CNS and pectoral fin buds. By contrast, tissues assigned to the crystalline archetype exhibited nuclei arranged in a low-density, anisotropic and regular fashion and included cells of the muscles, skin and YSL (Fig. 2 d,e). This bimodality becomes especially apparent when the archetypes are projected onto the embedding of the organisational feature space, with each covering an opposing hemisphere (Fig. 2 c). Moreover, we found that this bipartitioning was not a feature of a single developmental stage but could be identified throughout 12 and 72 hpf (Suppl. Fig. 9). These results show that the bimodal organisation of bona-fide tissues identified in the previous section (Fig. 1 f) extends to the whole organism and that local cellular arrangement follows one of two general organisational archetypes.

### nuQLOUD analysis links cadherin expression domains and organisational archetypes

To identify potential genetic regulators of this organisational bipartitioning into archetypes, we investigated the cadherin family of cell adhesion receptors, important regulators of tissue integrity and differential adhesion-based cell sorting^19,27^. While much is known about how cadherins function at the molecular, cellular, and biophysical level, their impact on tissue organisational diversity of whole organisms is less clear. To examine whether cadherins play a role in this context, we probed the relationship between specific cadherin expression and architecture by projecting *in toto* expression patterns of twelve highly expressed cadherins (Suppl. Fig. 10 a), defined by HCR, onto the nuQLOUD framework (Fig. 3 a). This revealed that each cadherin expression domain localised to discrete, albeit overlapping regions in feature space, indicating that cells expressing each specific cadherin had a range of organisation that is definable with our approach (Fig. 3 b). Clustering based on organisational similarity allowed cadherin expression domains to be grouped into two classes that mapped to the previously identified organisational archetypes (Fig. 3 c-e). This clustering of cadherin family members based on the organisation of expressing tissues was not predicted by similarities in their amino acid sequence or structures^28^ (Suppl. Fig. 11). Grouping cadherin expressing cells based on their transcriptional similarity as defined by single cell RNA sequencing^29^ generated a subdivision of cadherin family members into the same classes (Fig. 3 f, Suppl. Fig. 10 b,c). Pairwise co-expression analysis ruled out single-cell co-expression of multiple cadherins as reason for the organisational and transcriptional bimodal classification (Suppl. Fig. 10 d). Comparing the hierarchy of transcriptional and organisational similarity revealed that amorphously organised cadherin domains included more transcriptionally similar cells, while crystalline organised cadherin domains were more distinct in their gene expression profile, highlighting that diverse sets of gene expression may result in similar organisational patterns. The correlation between cadherin expression patterns and archetypal organisation suggested that individual cadherins may be required for the organisation of their respective expression domains. To test that, we systematically mutated individual cadherins using CRISPR/Cas9, a strategy that proved to effectively edit the respective loci as confirmed by sequencing (Suppl. Fig. 12 a,b). We specifically analysed the organisational effect on cells that normally would express the targeted cadherin, which we identified via HCR FISH (Suppl. Fig. 12 c). While cadherins are known to act redundantly^30^, loss of the amorphous class cadherins cdh2, cdh7a, cdh13 and cdh18a, resulted in their target tissues shifting detectably shifting towards the crystalline archetype (Suppl. Fig. 12 d-f). In addition to supporting functional roles for cadherin in promoting tissue archetypes, these analyses demonstrate how nuQLOUD can be used to highlight connections between transcriptional and tissue organisational patterns, links that can then be tested with perturbations using the same quantitative framework.

**Figure 3:**
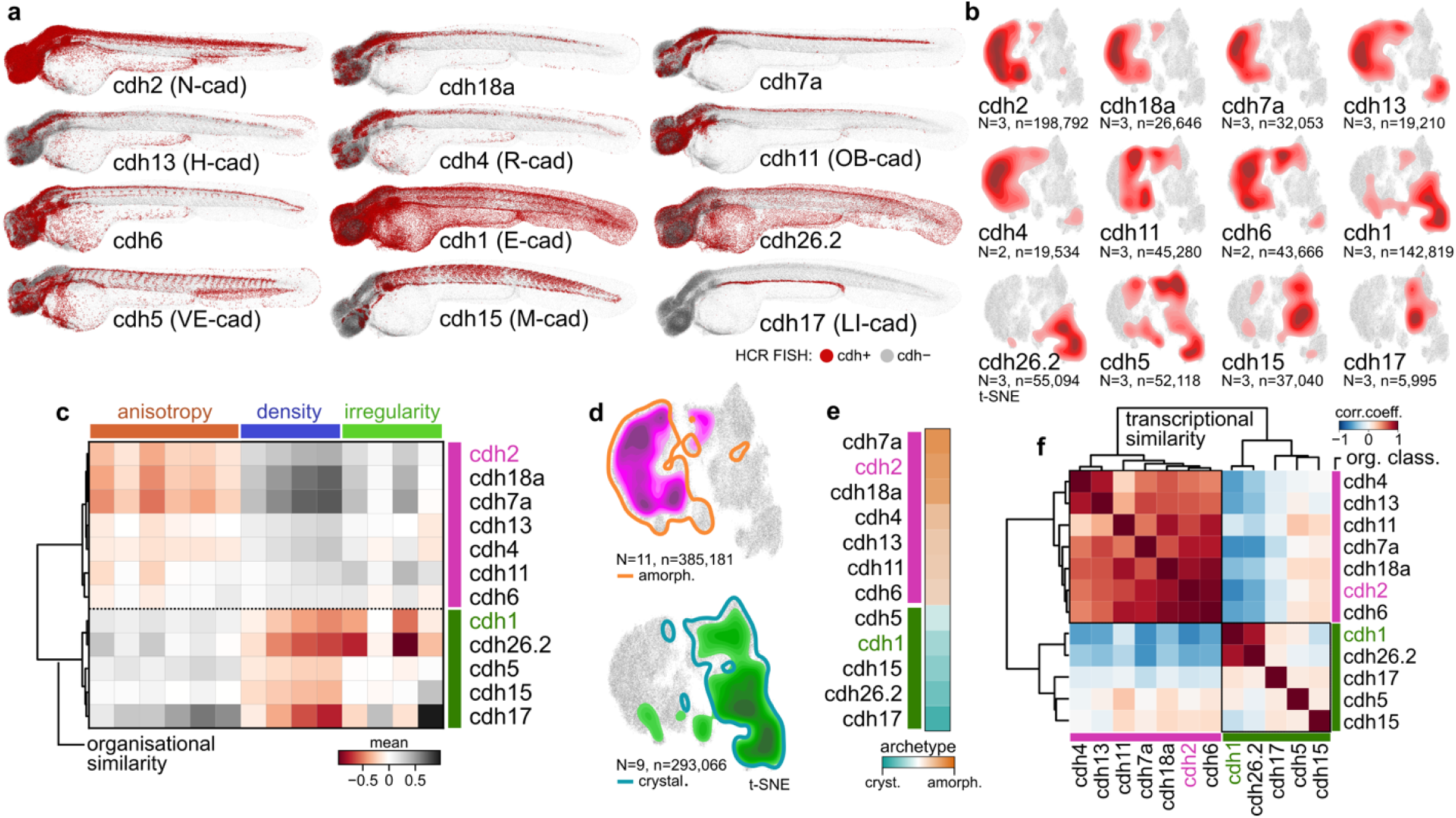
Organisational similarity identifies two classes of cadherin expressing cells that correlate with organisational archetypes. a: Cadherin expression domains are mapped on the point cloud representation; individual nuclei are classified as expressing/non-expressing based on fluorescence intensity of the label (HCR). Cadherins expressed in more than 1% of all cells were selected (Suppl. Fig. 10 a). b: Specific cadherin expression domains occupy distinct regions of organisational feature space. Domains either localise towards the left or right hemisphere. c: Clustering based on organisational features reveals two classes of cadherin expressing cells. Mean feature values of expression domains are z-scored. d: The two cadherin classes map to discrete halves of feature space and are predominantly contained by the amorphous and crystalline archetype respectively. e: The two cadherin classes map to organisational archetypes. Scale: percentage of archetype associating cells; cyan/orange 100% association, white 50% each. f: Auto-correlation clustering of cadherin expression domains based on single cell RNA sequencing data^29^ yields the same E- and N-cadherin-like classes as the classification based on organisational similarity. Pearson correlation coefficient. N: number of samples, n: total number of cells, dendrograms constructed from hierarchical agglomerative clustering using Ward linkage and Euclidean distance.

### N-Cadherin expression dynamics regulate archetype-switching during muscle development

To investigate the link between cadherin expression and organisational archetypes further, we dynamically analysed the tissue organisation of cadherin expression domains, reasoning that, if cadherins regulate organisation, they should maintain their organisational profile over time. We therefore selected E- and N-cadherin (cdh1, cdh2, E-/N-Cad) as representatives of the amorphous and crystalline organising cadherin classes and tracked their expression patterns using transgenic reporters (TgCRISPR(cdh1-mLanYFP)^31^, TgBAC(cdh2:cdh2-GFP))^32^ between 12 and 72 hpf (Fig. 4 a). E-cad expression remains high in the epidermis and other epithelial cells that maintain a crystalline organisation throughout. By contrast, the pattern of N-cad expression is dynamic, being expressed in most embryonic

**Figure 4:**
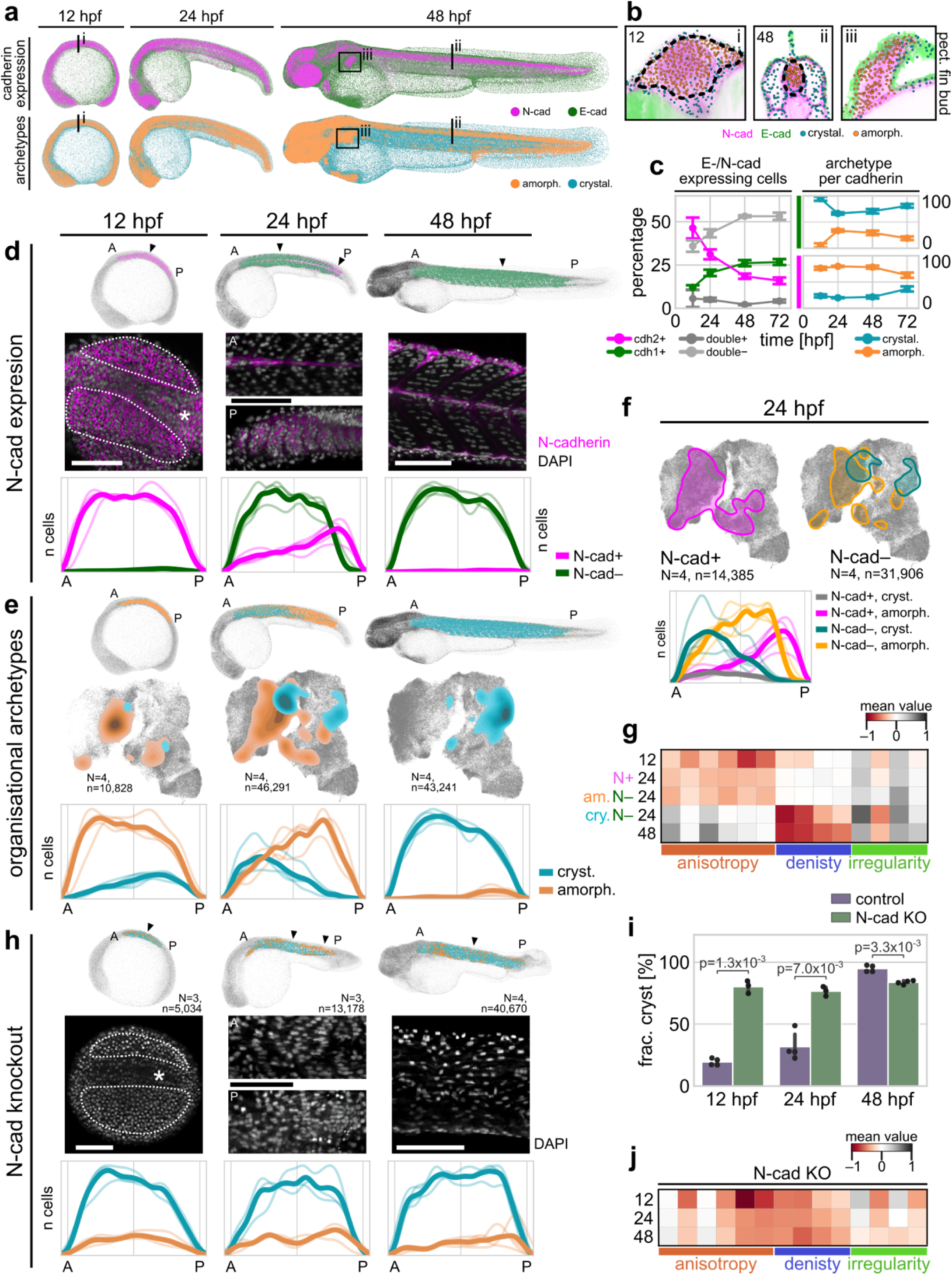
Changes in N-cad expression are linked to an amorphous to crystalline transition during muscle development. a, b: Expression patterns of E- and N-cad qualitatively match crystalline and amorphous archetypes. Grey dots: non-expressing cells; double-positive cells not shown. c: Association of cdh1/2 with crystalline/amorphous organisation is maintained throughout early development. N=4 samples/time point. P-value of amorphous vs. crystalline population per time point < 1.02×10-5 (cdh1), < 3.00×10-4 (cdh2); Welch’s t-test. d: Muscles downregulate N-cadherin expression in an anteroposterior (A-P) fashion. Arrowheads: position of representative images, illustrating downregulation of N-cadherin and organisational change. Dotted lines: muscle progenitors at 12 hpf; asterisk: notochord for orientation. Line profiles: number of cadherin expressing cells along A-P axis. 12 hpf: dorsal view, 24, 48 hpf: sagittal view. Ordinates are not normalised between time points. e: Organisational archetypes switch in an anteroposterior fashion. t-SNE and A-P line profiles illustrate organisational transition of muscle cells during development. f: Crystalline muscle cells have downregulated N-cadherin, while N-cadherin positive cells are organised amorphously during transition. g: Developing muscles become less isotropic and less dense when switching organisational archetype. h-i: Knockout (KO) of N-cadherin induces increase in crystalline organisation and loss of A-P pattern at 24 hpf. Transient KO of N-cadherin via CRISPR/Cas9 in F1. h: Representative images illustrate organisational effect of N-cadherin KO and line profiles show loss of A-P archetype-transition. i: Significantly more cells organise crystalline at 12 and 48 hpf in the KO than in WT control (Welch’s t-test). j: N-cadherin deficient cells organise at similar density as muscles at 48 hpf in WT but exhibit more heterogeneous isotropy features. All scale bars 100 µm. Solid lines in A-P line plots indicate mean cell number, smaller lines show individual sample’s profiles.

cells at 12 hpf, before becoming focussed to cells of the CNS and a subset of other tissues from 48 hpf onwards (Fig. 4 a-c, Suppl. Fig. 13). Interestingly, N-cad positive cells were assigned to the amorphous archetype across the analysed time window; an association that was agnostic to cell type. For example, mesenchymal cells of the pectoral fins and neurons of the brain have little commonality in lineage or function, despite sharing N-cad expression (Fig. 4 b). This dynamic link between E-/N-cad expression and archetypal tissue organisation further supports the idea that specific cadherin expression regulates local tissue organisation.

To test this in more detail, we next addressed this relationship within a single tissue, focussing on the developing fast-twitch muscles between 12 and 48 hpf, a well-studied morphogenic system that exhibits dynamically changing cadherin expression, where N-cad is expressed homogeneously early before becoming restricted to a superficial layer of slow muscle cells later^33^. We sub-selected developing muscles based on their nuclear morphology and tracked their cadherin expression at key time points, which confirmed that muscle cells inside each forming somite downregulate N-cad expression in an anteroposterior ‘wave-like’ manner (Fig. 4 d, Suppl. Movies 1-3). nuQLOUD analysis revealed that fast twitch muscle progenitors switch their organisational archetype from amorphous at 12 hpf to crystalline at 48 hpf (Fig. 4 e). At 24 hpf this organisational transition shows a similar pattern to N-Cad expression within the same tissue: posterior cells expressed N-cad and remained in an amorphous organisation whereas anterior cells were N-cad negative and showed crystalline organisation (Fig. 4 e). In the transition zone midway along the anteroposterior axis we identified a population of N-cad negative, amorphously organised cells, but no N-cad positive crystalline cells (Fig. 4 f), suggesting that cells downregulate their N-cad expression prior to their archetypal transition. Focusing on the organisational features over time revealed that muscle cells transition from an relatively unorganised, isotropic, and dense arrangement to a highly organised, anisotropic, and low-density configuration as N-cad expression decreased. This shift reflects their gradual alignment into parallel fibres, where the directional organisation of the fibres contributes to their anisotropic footprint (Fig. 4 g). The N-cad negative amorphous/crystalline populations at 24 hpf exhibited almost identical feature profiles as earlier/later time points, suggesting that the transition between archetypes was rapid. These findings are consistent with a model where N-cad expression locks cells in the amorphous archetype, an idea we tested directly by depleting N-cad function using CRISPR/Cas9 (Suppl. Fig 14 a-e). Early myotomes of N-cad knockout (KO) embryos exhibited predominantly crystalline organisation already from 12 hpf onward and throughout the observed time window (Fig. 4 e,f). N-cad deficient cells maintained a high degree of anisotropic organisation, yet exhibit reduced density throughout (Fig. 4 h,g). Taken together, these results show that N-cad expression is required for the maintenance of amorphous organisation of early somites.

### Tissues undergo amorphous to crystalline switch in the absence of N-cad function

The finding that downregulation of N-cad expression enables tissues to transition from an amorphous to a more crystalline organisation predicts that the continued high-level expression of N-cad by tissues such as the CNS could be required to maintain their amorphous archetype. To test this hypothesis, we expanded our analysis of N-cad KO to whole embryos between 12 and 48 hpf, starting at a developmental stage, when N-cad has just reached its full expression (Suppl. Fig 15 a). 12 hpf saw a significant reduction in the fraction of cells showing amorphous organisation with an increase in crystalline organisation (Fig. 5 a, b). Moreover, N-cad KO embryos at this stage did not exhibit a significant reduction in total cell count over WT, indicating that the identified change in organisation isn’t a consequence of defects in cell proliferation or viability. However, at later stages N-cad KO embryos showed the previously described morphological phenotypes and lower cell numbers overall (Fig. 5 c, Suppl. Fig 15 b-c), meaning that changes in tissue organisation at these stages could be a secondary consequence of more global developmental defects. To study the organisational effects downstream of N-cad loss more directly, we focussed on the CNS, the major N-cad expressing tissue at 48 hpf. We generated identifiable N-cad deficient clones by combining a genetic mosaic approach with transgenic reporters for N-cad expression and neural identity. Mosaic knockout (mKO) of N-cad was achieved by injecting Cas9 + sgRNAs against N-cad into a single cell at the 8-cell stage (Fig. 5 d). This significantly reduced the general morphological defects observed in 1-cell stage CRISPR injections or N-cad homozygous mutants lines (Suppl. Fig. 14 f). To identify N-cad deficient clones, we took advantage of the fact that most cells of the CNS at 48 hpf coexpress N-cad and the neuronal reporter Neural Beta-Tubulin (NBT:dsRed) (Fig. 5 e). Therefore, we identify CRISPANT cells (i.e., cells that would express N-cad normally, but have lost gene function) as cells that express NBT but not N-cad in the mKO condition (Fig. 5 d, e). We found a significant reduction of such double positive cells in mKO samples over WT (Fig. 5 e), confirming the efficacy of the approach. Using whole-brain light sheet microscopy (Fig. 5 f, Suppl. Fig. 16), we quantitatively compared the organisation of N-cad deficient (NBT+/Cdh2-GFP-) and surrounding WT neurons (NBT+/Cdh2-GFP+) in mKO embryos at 48 hpf. In WT animals, analysed NBT+ cells organised amorphously, as did NBT+/Cdh2-GFP+ cells in the mKO conditions (Fig 5 g-i). By contrast, NBT+/Cdh2-GFP-neurons within the same sample were assigned at a significantly higher frequency to the crystalline archetype (Fig. 5 g, h). The archetypal switch observed after N-cad loss was due to a decrease in cell density and an increase in regularity (Fig. 5 i). These results show that N-cad expression is not only required for the maintenance of amorphous organisation during muscle development, but serves the same role during the development of the CNS. Together with the dynamic and phenotypic findings from *in toto* analysis, this supports a model where N-cad is a general driver of the amorphous archetype during zebrafish development.

**Figure 5:**
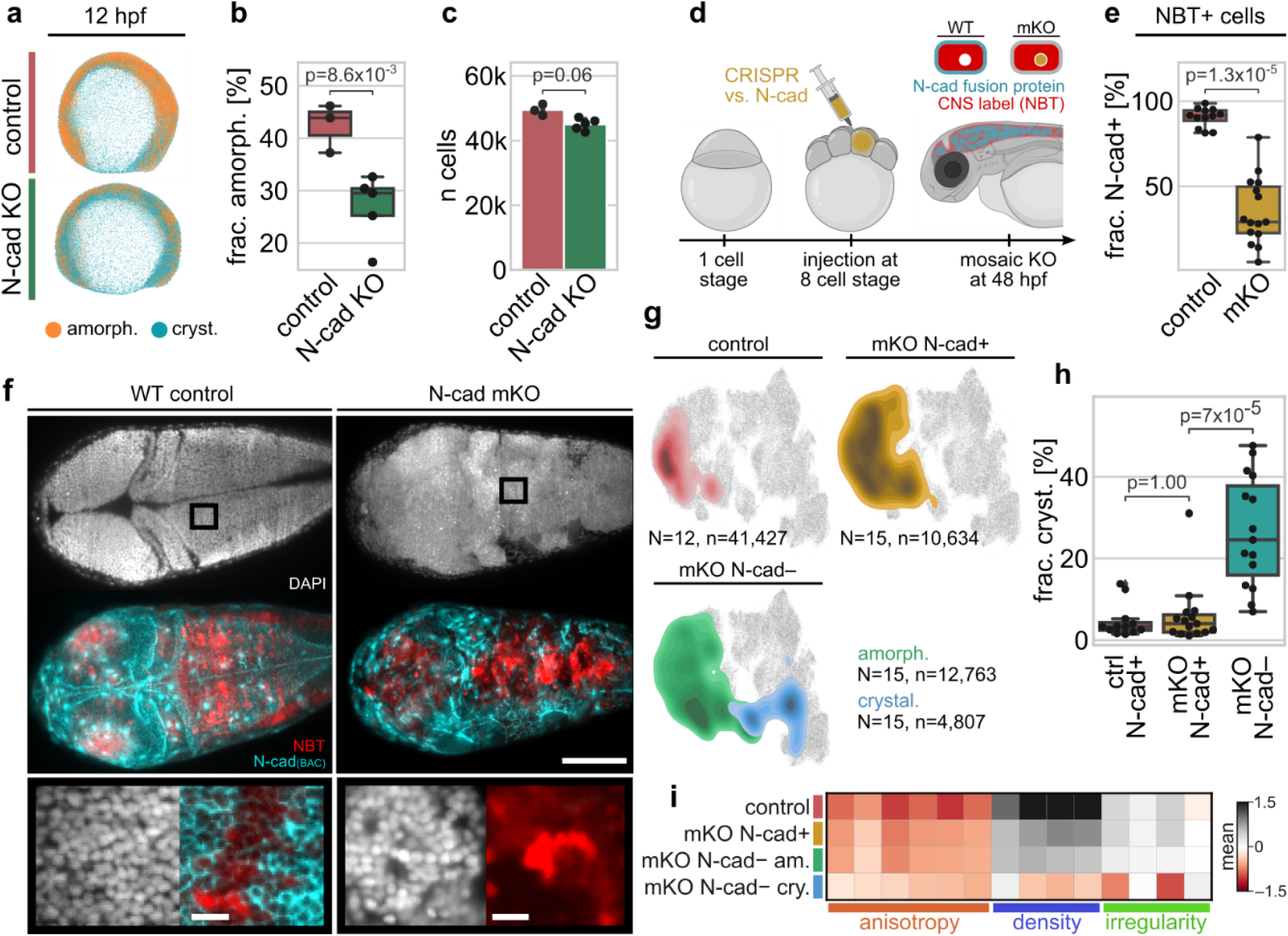
Clonal N-cad loss induces localised organisational shift towards crystalline archetype. a-c: Knockout of N-cadherin decreases frequency of amorphous organisation at 12 hpf. a: Representative renderings of uninjected (control) and N-cadherin KO embryos; 12 hpf, nuclei segmented and coloured according to archetype. KO was achieved via injection of Cas9 + sgRNAs against N-cadherin at the 1-cell stage of embryonic development. Fraction of cells organising amorphously decreases about 30% between control and KO (b) while the number of cells between the conditions did not change significantly (c). d: Mosaic KO (mKO) of N-cadherin is achieved by injecting Cas9 + sgRNAs into a single cell at the 8-cell stage of embryonic development. Manipulated cells are identified via combinatorial labelling: cells of the CNS co-express NBT and N-cadherin in WT conditions; mKO cells have lost N-cadherin and only express NBT. e: mKO is efficient and reduces the fraction of NBT+ cells that are N-cadherin+ from ∼98% to ∼34%. f: Whole-brain light sheet microscopy confirms morphological phenotype and indicates alteration in tissue organisation in mKO clones. Scale bars: overview 200 µm, zoom-in 15 µm. g: Projection of NBT+ cells onto organisational feature space representation reveals archetypal shift towards crystalline organisation exclusive in N-cadherin mKO cells. Organisation of segmented nuclei was quantified per sample and subsequently aligned with the in toto organisational feature space. N: number of samples, n: total number of analysed cells. h: Fraction of crystalline organised cells increases significantly in N-cad mKO cells over N-cad+ cells; the latter organise not significantly different from N-cad+ cells of control embryos. i: Shift towards crystalline organisation is driven by a decrease in cell density and an increase in regularity. Indicated p-values were calculated via Welch’s t-test.

## Discussion

Advances in light microscopy and transcriptomics have led to an explosion of interest in tissue heterogeneity in a wide variety of experimental systems from organoids^34^ to synthetic embryos^19^, and whole small organisms^13^. Progress towards a more general mechanistic understanding of tissue diversification depends on integrating data collected from different contexts. The motivation behind *nuQLOUD* was to provide a common imaging-based framework that is flexible and general enough to encompass the high organisational variation that is found between tissues and organisms. We achieved this goal by sacrificing cell-scale morphological features, which are more challenging to quantify and are better-suited for characterising cell-type differences, and instead focussing on how cells are arranged in 3D. This focus on the organisation of tissues rather than cells is analogous to how architecture generally focuses on the layout of buildings rather than bricks. Reducing embryonic complexity to the positions of all cell nuclei, achieved using established in toto segmentation methods^10^, allowed us to leverage the computational efficiency of point-cloud-based frameworks and analyse large numbers of embryos each comprising >100,000 cells. We demonstrate that this can be done by simply adding DAPI, a universal DNA stain that removes the need for specific genetic reporters, thus nuQLOUD can be used to quantify tissue architecture in basically any biological specimen, essentially *gratis*.

One goal of the nuQLOUD approach was to provide a novel way to define tissue organisation at the whole organism scale. General descriptors of tissue state, such as the epithelial-mesenchymal paradigm, are extremely powerful as they enable the transfer of concepts and mechanisms between a variety of biological contexts^35^. While mesenchymal and epithelial states are normally identified using diagnostic markers, they were originally defined via their distinct morphological and organisational characteristics^36^. Such architectural features, often obvious to the trained eye, are challenging to quantify and define systematically. Addressing exactly this challenge, nuQLOUD shows that embryonic complexity can be reduced to crystalline and amorphous tissue archetypes that appear analogous to epithelia and mesenchyme, respectively. However, it’s clear that these tissue archetypes are not simply ‘organisational proxies’ of classical mesenchymal and epithelial states. For example, while the crystalline archetype comprises epithelia such as skin and pronephros, it also includes non-epithelial tissues like muscles. Likewise, the amorphous archetype includes pectoral fin mesenchyme but also branchial arches, spinal cord neurons and eyes. These tissues are amorphous, as they are denser and less positionally ordered than epithelia, but they don’t fit the definition of loosely organised bona fide mesenchyme. Indeed, nuQLOUD’s local organisational features select for cell collectives and tissues rather than individual mesenchymal cells. Thus, the amorphous archetype describes cells that are in-between epithelia and mesenchyme on the organisational spectrum. Of relevance here is the emerging concept that such ‘E/M hybrid’ states, also termed partial EMT, correlate with increased fate plasticity or ‘stemness’ in contexts such as cancer and reprogramming^37^. Intriguingly, time-resolved analysis revealed that most internal cells in early embryos are initially amorphous before undergoing a progressive ‘crystallisation’ that correlates with organ progenitor assembly. Such ’amorphous to crystalline transitions’ again appear analogous to mesenchymal to epithelial transitions (MET), and the related jamming transition^21,38^, however they can be identified systematically from changes in cell arrangement alone. Combined, these data support the conclusion that organ maturation displays a general organisational signature that can be reliably detected using the nuQLOUD framework, as demonstrated for example in the maturation of fast twitch muscles.

The quantitative readout of tissue organisation provided by nuQLOUD allowed us to explore the relationship between cadherin expression and organisational archetypes. Integrating spatial expression data of 12 major cadherins into the nuQLOUD framework revealed two distinct groups: cadherins linked to crystalline tissues, such as E-cad, and those associated with amorphous tissues, including N-cad. Time-resolved analysis revealed that N-cad expression is tightly correlated with the amorphous archetype, being expressed by the majority of cells at early stages and becoming replaced by more tissue-specific cadherins as cells undergo amorphous to crystalline transitions. By contrast, cells of the nervous system displayed high N-cad expression and amorphous archetype organisation across all stages studied, consistent with its described role in nervous development. N-cad depletion, either through normal developmental downregulation or targeted perturbations, led to cells switching their organisation from amorphous to crystalline, identifying N-cad as a central regulator of the amorphous archetype. These data support a model where N-cad downregulation and exit from the amorphous archetype represents a common gating mechanism regulating non-neural tissue assembly. Interestingly, recent findings show that inactivation of N-cad function increases the self-organisation potential of cultured mammalian gastruloids^40^, consistent with the proposal that regulation of N-cad expression may play a similar gating function during the formation of synthetic embryos. It will be very interesting to explore whether archetypal organisational patterns and transitions are also a predictive feature tissue assembly in such synthetic embryo systems. Looking ahead, we can imagine a number of ways in which nuQLOUD be combined with other approaches to address key questions in tissue biology. Of general interest is how changes in the cell arrangement feed back on signalling pathways that control cell fate, whether through pathways regulated via cell-cell coupling, such as Hippo and Notch, or by changing the reach and distribution of morphogens. Recent work on organ development^41^, tissue patterning^42^, and immune regulation^43^ has highlighted that differences in tissue organisation can be as influential in cellular decision making as signalling pathways. Developing a standard language for the quantification of cellular arrangement, like the one presented here, will be instrumental for a more integrated understanding of the interplay between genetic and morphological factors that enable contextual decision-making in physiology and disease.

The current nuQLOUD framework is not without limitations. First, it assumes continuous arrangement over a characteristic length scale, typically one nearest neighbour on the Delaunay graph. While this assumption is appropriate for analysing tissues with dimensions exceeding ∼30 µm, it is less optimal for narrow structures, such as blood vessels^44^, or isolated cells, such as macrophages and fibroblasts, which are sampled along with their surrounding tissue. Second, organisational features in nuQLOUD are defined as local averages, which may average out tissues that exhibit different structural properties along different axes. For example, at later stages the retina shows regular, layered organisation in one direction, while being more amorphous in orthogonal planes. To account for this, future versions of nuQLOUD will include directional organisational features, such as directed variation of cell density. Our investigation of cadherins identified N-cad as a key player in regulating amorphous organisation, however the lack of phenotype resulting from inactivation of others does not mean that they play no role, as cadherin are known to act redundantly.

## Methods

### Zebrafish handling

Zebrafish (Danio rerio) strains were maintained, grown and bred following the standard procedures described in^45^. All experiments were conducted in accordance with the regulation and guidelines of the veterinary office of the University of Zürich and the Canton of Zürich, Switzerland. Embryos were staged following^25^.

### Chemical treatment

Embryos were treated with 0.002% N-phenylthiourea (PTU, Sigma-Aldrich) from 24 hpf on to prevent pigmentation. For immobilisation of embryos older than 24 hpf during screening they were treated with 0.01% tricaine methanesulfonate (MS222, Sigma-Aldrich). Moreover, highly concentrated tricaine (300 mg/L) was used to euthanize embryos prior to fixation. To aid and accelerate dechorination of embryos between 24 and 48 hpf, they were treated with 0.05% pronase^46^.

### Transgenic fish lines

To visualise gene expression live and without staining, transgenic zebrafish lines were used. The nervous system was visualised using Tg(NBT:DsRed)^47^, E-cadherin expression was visualised using either TgCRISPR(cdh1-mLanYFP) or TgCRISPR(cdh1-tdTomato)^31^. N-cadherin expression was visualised using TgBAC(cdh2:cdh2-GFP)^32^.

### Fixation of samples and nuclear staining

Prior to imaging, zebrafish embryos at stages before 48 hpf were manually dechorionated using Dumont #5 forceps; embryos at later time points were already hatched. Embryos were euthanized using tricaine and fixed for one hour at room temperature in 4% paraformaldehyde (PFA) in 1X phosphate-buffered saline (PBS). Subsequently, the samples were rinsed three times with PBS Tween (PBS + 0.05% Tween-20, Thermo Fisher Scientific, PBS-T), permeabilized for one hour in PBS Triton (PBS + 0.1% Triton X-100, Thermo Fisher Scientific) and rinsed again with PBS-T. The nuclei of the samples were stained by treating the sample with 1X 4′,6-diamidino-2-phenylindole (DAPI) for two hours at room temperature. After staining, the samples were rinsed with PBS-T, stored in PBS-T at 4°C and imaged within 14 days.

### Fluorescent labelling of transcripts via HCR RNA FISH

To identify cells that expressed genes of interest, we labelled their transcripts using hybridisation chain reaction RNA fluorescent in situ hybridisation (HCR RNA FISH)^48^. Probes were designed by and ordered from Molecular Instruments. Fluorescent amplifiers with emission wavelengths of 594 nm (red) or 647 nm (infra-red) were used. Generally, two sets of HCR probes were multiplexed within individual samples to increase imaging throughput. To stain samples with HCR RNA FISH, embryos were fixed for one hour using 4% PFA. Subsequently, samples were washed for three cycles with PBS-T and permeabilised via a methanol dehydration sequence (25, 50, 75 % methanol in PBS-T for 15 min, 100% methanol for 1h). A reversed sequence was used to rehydrate samples with 10 min per step and four concluding wash cycles. Afterwards, samples were pre-hybridised using the probe hybridisation buffer at 37 °C for 30 min and HCR probes were applied (2 pmol per probe) for 12 – 16 h at 37 °C. To remove unbound probes, samples were incubated four times for 15 min in probe wash buffer and two times 5 min in 5x saline-sodium citrate + tween (ssct) buffer. Embryos were pre-amplified in amplification buffer for 30 min and then incubated with hairpin solution for 12 – 16 h. Hairpin solution was generated by mixing separately heat-shocked (30 s at 95 °C) amplifier-specific hairpin h1 and hairpin h2 solutions in amplification buffer. The staining was completed by a final five cycles of PBS-T washes. Samples were kept in PBS-T afterwards and either further processed or imaged.

### F0 CRISPR knockout and validation

F0 knockouts were generated via injection of a sgRNA/Cas9 mix into early-stage zebrafish embryos. sgRNAs sequences are summarised in (Suppl. Table 2) The injection mix was composed of Cas9-2xNLS protein (1 mg/µL), the sgRNAs (128 ng/µL) and phenol red (0.025%). Full knockouts were achieved by injecting 1-2 nL of injection mix into the yolk at the 1-cell stage of embryonic development, mosaic knockouts (mKO) were achieved by injecting into one cell at the 8-cell stage of development (∼1h 10min post fertilisation at 25°C). Cas9 protein was sourced from the Protein Expression and Purification Core Facility of the European Molecular Biology Laboratory (EMBL) and was kept in media containing Hepes, KCl, and Glycerol at -80 °C.

To verify the efficacy of sgRNAs, the homology regions of the used sgRNAs were amplified and sequenced to detect DNA sequence polymorphisms. For that, injected embryos were grown to 4 dpf and euthanized using tricane. Genomic DNA was extracted from individual samples using 40 µL QuickExtract DNA extraction solution (Lucigen QE0905T). Embryos were incubated for 15 min at 25 °C, 5 min at 65 °C and 2 min at 95 °C and vortexed in between incubation steps. Subsequently, solid debris was removed by centrifuging 1 min at 13,000 rcf. To amplify the region of sgRNA homology, flanking primer pairs were designed using the software tool Primer3^49^ with default parameters. Primer sequences are summarised in Suppl. Table 3. Polymerase chain reaction (PCR) was performed using Taq polymerase (NEB M0267S) and PCR products were analysed via gel electrophoresis. Band locations were predicted based on the size of the cut or uncut genomic DNA sequence and DNA was recovered via gel extraction and purification (QIAGEN 28604). Sanger sequencing was used to detect sequence polymorphisms which indicated sgRNA efficacy (Suppl. Fig. 12, 14).

### Microscopy

Samples were mounted in 1% low-melting agarose solution in 1X E3 fish embryo medium by aspirating them head-first into 20µL glass capillaries (BRAND Transferpettor caps, Merck) using Transferpettor piston rods (BRAND, Merck).

Light sheet microscopy was performed on a Zeiss Z.1 Lightsheet microscope. All images were acquired using a Zeiss W Plan Apochromat 20×1.0 corrected water immersion objective and a set of two Zeiss 10x illumination objectives. For *in toto* acquisition the detection objective was de-zoomed by factor 0.45; 0.65 for partial brain imaging. The beam waist of the light sheet was optimised for maximum uniformity across the field of view. Images were recorded using scientific Complementary Metal–Oxide–Semiconductor (sCMOS) cameras (PCO edge 5.5) with 1920×1920 pixels. During acquisition, the camera sensors were cropped in a portrait fashion (1200-1600×1920 pixels) to exclude non-signal areas, improving acquisition speed, and decreasing data volume.

For whole-organism acquisition, samples were imaged from four orthogonal sides starting with a lateral view with the ventral-dorsal axis of the sample going from left to right and continuing in steps of 90° clockwise along the anteroposterior axis of the sample. For 12 hpf samples one such imaging volume was sufficient to capture a whole sample. For more developed samples, several overlapping (10%-20%) four-view imaging volumes were acquired along the anteroposterior axis of the sample; 24 hpf: 2 volumes, 48 hpf: 4 volumes, 72 hpf: 5 volumes. The z-spacing of consecutive frames was 1µ and for each frame the left and right light sheet illumination was recorded separately. Acquisition is illustrated in Suppl. Fig. 1. During acquisition the ‘Pivot Scan’ option was turned on to reduce illumination artefacts. The light sheet offset was manually calibrated for each sample individually by contrast maximisation. Illumination settings were chosen to minimise acquisition time. To avoid photobleaching during setup, spatial alignment and multiview setup was performed using the DAPI channel. Other fluorescent channels were set up using single frame acquisition. Multicolour imaging was performed sequentially by prioritising z-stacks over colour, i.e., full z-stacks were acquired repeatedly with single colours. The whole imaging process can hence be summarised as: light sheet direction → z-stack → colour → volume → tiles.

### Multi-view fusion of light sheet microscopy data

Raw light sheet images were stored as czi (Zeiss) files. The MVRegFus package (available at https://github.com/m-albert/MVRegFus) was used to fuse the acquired data into one image volume with isotropic resolution and locally optimised quality. First, the opposing light sheet illuminations were fused per frame maximising the normalised discrete cosine transformed Shannon Entropy (DCTS) to select the highest contrast elements for the fused image. Subsequently, overlapping image volumes were pairwise fused, again using DCTS to select the highest contrast regions. The order of fusion was such that orthogonal views were fused in clockwise fashion along the anteroposterior axis. Frames along the anteroposterior axis were connected by fusing overlapping volumes acquired with the same rotational angle. The final fusion was performed using Lucy-Richardson deconvolution. The brightness of individual frames was normalised using Contrast Limited Adaptive Histogram Equalization. For processing, voxel intensities were thresholded at 200 AU, to circumvent fusion artefacts caused by thermal noise. Images were binned 4×4×1 (x,y,z) during processing to increase performance. The final isotropic resolution of the fused image was 1 µm/voxel. The processed images were stored in the hdf5 derived ims format for efficient compression and handling. The fusion pipeline is illustrated in Suppl. Fig. 1.

### Nuclear segmentation via TGMM

Individual nuclei were segmented in 3D from volumetric microscopy images using the Tracking with Gaussian Mixture Models (TGMM 2.0) software^10^. The background of individual images was estimated by measuring the maximum intensity of regions not containing nuclear signal using the image analysis software Fiji^50^. The anisotropy parameter was set to 1.0 since the processed images had isotropic resolution. The minimal and maximal size of detected nuclei was determined by measuring nuclear sizes in fluorescence images and was set to 150 and 4000 voxels respectively. To suppress mis-segmentations, the covariance matrix of the 3D Gaussian distribution was regularised such that its eigenvalues were bounded between 0.02 and 0.1, limiting the size of the three principal axis of the distribution. Additionally, total eccentricity was limited to ɛ = 9. All other parameters were either left at default values or they were empirically determined to their used values. All parameters are summarised in Suppl. Table 1. The TGMM software was implemented in C++ and was released with a command line-based interface or a rudimentary graphical user interface. Both were not suitable for automated segmentation of multiple images in parallel and generally struggled with processing more than a single image at a time. We therefore developed a python-based API which managed the tool’s parameters, and the image- and segmentation file paths. Moreover, it enabled the parallel processing of multiple images simultaneously and automatically transformed the TGMM output files into a DataFrame representation. The software is publicly available on https://github.com/max-brambach/tgmm_utility.

### Generation of Voronoi Diagrams

Three-dimensional Voronoi diagrams were generated using a modified version of the software voro++^51^. We modified the software to enable the generation of a radially restricted Voronoi diagram using command line input, which enabled the integration of the tool into a python-based workflow. The radial restriction was achieved by initialising every Voronoi cell as a dodecahedron with edge length 100 voxels (≙ µm). Overlapping Voronoi cells were subsequently cropped to a regular Voronoi diagram. Boundary cells were limited to their initial shape and could therefore be identified by consisting of faces that were not shared with other Voronoi cells. The modified voro++ version is publicly available on https://github.com/max-brambach/voro. To suppress boundary effects on the Voronoi diagram, we developed a method to restrict the size and shape of boundary cells based on the distribution of their neighbouring cells using auxiliary points and a secondary Voronoi diagram (Suppl. Note 1)

### Single cell RNA sequencing data processing

Single cell RNA sequencing data of early zebrafish development was obtained from ^29^ and processed using scanpy. Pre-processing was performed following ^52^. Initially, cells were filtered out that expressed less than 200 genes and genes were removed that were expressed in less than 3 cells. The numbers of counts per cell were normalised to 10,000 and logarithmized. Highly variable genes were identified following ^53^ with minimum mean 0.125, maximum mean 3, and minimum dispersion 0.5. Individual genes were scaled to unit variance and maximum standard deviation was limited to 10. Differential gene expression analysis was performed according to ^54^ using the Mann-Whitney U-test.

### Statistical analysis and data representation

If not stated otherwise, statistical significance was tested using non-parametric tests. For unpaired samples, the Mann-Whitney U test was used and for paired samples the Wilcoxon signed-rank test was used. If not stated otherwise, N denotes the number of samples (replicates) and n the number of cells (observations). For statistical analysis, observations are grouped by sample.

Box plots illustrate data distributions by showing their median (central line), interquartile range (box), 1.5 x interquartile range (whiskers), and outliers (flyers). Error bars on bar plots indicate standard deviation of the mean if not stated otherwise.

For low dimensional embedding either t-SNE or UMAP were used. t-SNE representations were generated following ^55^ using principal component initialisation, dual perplexities (50 and 500), learning rate n/12 and the cosine distance. 25,000 randomly selected observations were pre-embedded using 250 iterations at exaggeration 12, and momentum 0.5, followed by 750 iterations at exaggeration 1, and momentum 0.8. The remaining dataset was mapped onto the pre-embedding using a k-nearest neighbour search in feature space. Finally, the full embedding was optimised with perplexity 30, 500 early exaggeration iterations (exaggeration 4, momentum 0.5) and 500 iterations at exaggeration 3, and momentum 8. UMAP embeddings were generated using minimum distance 0.5, spread 1.0, alpha 1.0, gamma 1.0, negative sample rate 5 and initial spectral embedding.

## Data and code availability

All data supporting the findings of this study are available within this article and its supplementary files or from the corresponding authors upon reasonable request. All raw light sheet data (post multi-view fusion) is available on the BioImage Archive (https://www.ebi.ac.uk/bioimage-archive/), accession number S-BIAD1405. A python implementation of nuQLOUD is available at https://github.com/max-brambach/nuQLOUD. A modified version of voro++ accompanying nuQLOUD is available at https://github.com/max-brambach/voro. Tools to efficiently run nuclear segmentation using TGMM are available at https://github.com/max-brambach/tgmm_utility. The multi-view fusion algorithm MVRegFus is available at https://github.com/m-albert/MVRegFus.

## Author contributions

MB and DG conceived the project and designed experiments. MB performed all experiments, performed light sheet microscopy, analysed images, developed the nuQLOUD software, performed data analysis. MA developed multi-view fusion software. MB and JJ performed F0 CRISPR KOs. RB provided transgenic zebrafish lines and optimised HCR staining. MB and DG interpreted the data and wrote the manuscript with input from all authors.

## Acknowledgements

Francesca Peri and lab, François Nédélec, Damian Brunner, Jana Wittmann and all members of the Gilmour lab for discussion. Cornelia Henkel for fish care. Urs Ziegler, Jana Döhner, Joana Delgado Martins of the UZH Center for Microscopy and Image Analysis for support with and infrastructure for image acquisition and analysis. Kim Remans and the Protein Expression and Purification Core Facility of the European Molecular Biology Laboratory for Cas9. Mark Cronan and David Tobin for E-Cad reporter lines. SNSF Grant 176235 to DG and UZH Forschungskredit FK-21-084 to MB.

## References

1. Weinreb, C., Rodriguez-Fraticelli, A., Camargo, F. D. & Klein, A. M. Lineage tracing on transcriptional landscapes links state to fate during differentiation. Science 367, eaaw3381 (2020).

2. Farrell, J. A. et al. Single-cell reconstruction of developmental trajectories during zebrafish embryogenesis. Science 360, eaar3131 (2018).

3. Spanjaard, B. et al. Simultaneous lineage tracing and cell-type identification using CRISPR–Cas9-induced genetic scars. Nat. Biotechnol. 36, 469–473 (2018).

4. Thompson, D. W. On Growth and Form. (Cambridge University Press, Cambridge, UK, 1917).

5. Domcke, S. et al. A human cell atlas of fetal chromatin accessibility. Science 370, eaba7612 (2020).

6. Cao, J. et al. A human cell atlas of fetal gene expression. Science 370, eaba7721 (2020).

7. Huisken, J., Swoger, J., Del Bene, F., Wittbrodt, J. & Stelzer, E. H. K. Optical Sectioning Deep Inside Live Embryos by Selective Plane Illumination Microscopy. Science 305, 1007–1009 (2004).

8. Keller, P. J., Schmidt, A. D., Wittbrodt, J. & Stelzer, E. H. K. Reconstruction of Zebrafish Early Embryonic Development by Scanned Light Sheet Microscopy. Science 322, 1065–1069 (2008).

9. Shah, G. et al. Multi-scale imaging and analysis identify pan-embryo cell dynamics of germlayer formation in zebrafish. Nat. Commun. 10, 5753 (2019).

10. McDole, K., et al. In Toto Imaging and Reconstruction of Post-Implantation Mouse Development at the Single-Cell Level. Cell 175, 859–876.e33 (2018).

11. Driscoll, M. K. et al. Robust and automated detection of subcellular morphological motifs in 3D microscopy images. Nat. Methods 16, 1037–1044 (2019).

12. Stringer, C., Wang, T., Michaelos, M. & Pachitariu, M. Cellpose: a generalist algorithm for cellular segmentation. Nat. Methods 18, 100–106 (2021).

13. Vergara, H. M. et al. Whole-body integration of gene expression and single-cell morphology. Cell 184, 4819–4837.e22 (2021).

14. Farhadifar, R., Röper, J.-C., Aigouy, B., Eaton, S. & Jülicher, F. The Influence of Cell Mechanics, Cell-Cell Interactions, and Proliferation on Epithelial Packing. Curr. Biol. 17, 2095–2104 (2007).

15. Garner, D. et al. Connectomic reconstruction predicts visual features used for navigation. Nature 634, 181–190 (2024).

16. Kintner, C. Regulation of embryonic cell adhesion by the cadherin cytoplasmic domain. Cell 69, 225–236 (1992).

17. Takeichi, M. Cadherin Cell Adhesion Receptors as a Morphogenetic Regulator. Science 251, 1451–1455 (1991).

18. Nose, A., Nagafuchi, A. & Takeichi, M. Expressed recombinant cadherins mediate cell sorting in model systems. Cell 54, 993–1001 (1988).

19. Bao, M. et al. Stem cell-derived synthetic embryos self-assemble by exploiting cadherin codes and cortical tension. Nat. Cell Biol. 24, 1341–1349 (2022).

20. McMillen, P., Chatti, V., Jülich, D. & Holley, S. A. A Sawtooth Pattern of Cadherin 2 Stability Mechanically Regulates Somite Morphogenesis. Curr. Biol. 26, 542–549 (2016).

21. Mongera, A. et al. A fluid-to-solid jamming transition underlies vertebrate body axis elongation. Nature 561, 401–405 (2018).

22. Tsai, T. Y.-C. et al. An adhesion code ensures robust pattern formation during tissue morphogenesis. Science 370, 113–116 (2020).

23. Hulpiau, P. & van Roy, F. New Insights into the Evolution of Metazoan Cadherins. Mol. Biol. Evol. 28, 647–657 (2011).

24. Nichols, S. A., Roberts, B. W., Richter, D. J., Fairclough, S. R. & King, N. Origin of metazoan cadherin diversity and the antiquity of the classical cadherin/β-catenin complex. Proc. Natl. Acad. Sci. 109, 13046–13051 (2012).

25. Kimmel, C. B., Ballard, W. W., Kimmel, S. R., Ullmann, B. & Schilling, T. F. Stages of embryonic development of the zebrafish. Dev. Dyn. 203, 253–310 (1995).

26. Cislo, D. J. et al. Active cell divisions generate fourfold orientationally ordered phase in living tissue. Nat. Phys. 19, 1201–1210 (2023).

27. Maître, J.-L. et al. Adhesion Functions in Cell Sorting by Mechanically Coupling the Cortices of Adhering Cells. Science 338, 253–256 (2012).

28. Nollet, F., Kools, P. & van Roy, F. Phylogenetic analysis of the cadherin superfamily allows identification of six major subfamilies besides several solitary members1. J. Mol. Biol. 299, 551–572 (2000).

29. Farnsworth, D. R., Saunders, L. M. & Miller, A. C. A single-cell transcriptome atlas for zebrafish development. Dev. Biol. 459, 100–108 (2020).

30. Astick, M., Tubby, K., Mubarak, W. M., Guthrie, S. & Price, S. R. Central Topography of Cranial Motor Nuclei Controlled by Differential Cadherin Expression. Curr. Biol. 24, 2541–2547 (2014).

31. Cronan, M. R. & Tobin, D. M. Endogenous Tagging at the cdh1 Locus for Live Visualization of E-Cadherin Dynamics. Zebrafish 16, 324–325 (2019).

32. Revenu, C. et al. Quantitative cell polarity imaging defines leader-to-follower transitions during collective migration and the key role of microtubule-dependent adherens junction formation. Development 141, 1282–1291 (2014).

33. Cortés, F. et al. Cadherin-Mediated Differential Cell Adhesion Controls Slow Muscle Cell Migration in the Developing Zebrafish Myotome. Dev. Cell 5, 865–876 (2003).

34. Wahle, P. et al. Multimodal spatiotemporal phenotyping of human retinal organoid development. Nat. Biotechnol. 41, 1765–1775 (2023).

35. Dongre, A. & Weinberg, R. A. New insights into the mechanisms of epithelial–mesenchymal transition and implications for cancer. Nat. Rev. Mol. Cell Biol. 20, 69–84 (2019).

36. Hay, E., Fleischmajer, R. & Billingham, R. Epithelial-mesenchymal interactions. in Proceedings of the 18th Hahnemann Symposium. Philadelphia, PA: Williams and Wilkins (1968).

37. Grande, M. T. et al. Snail1-induced partial epithelial-to-mesenchymal transition drives renal fibrosis in mice and can be targeted to reverse established disease. Nat. Med. 21, 989–997 (2015).

38. Petridou, N. I., Corominas-Murtra, B., Heisenberg, C.-P. & Hannezo, E. Rigidity percolation uncovers a structural basis for embryonic tissue phase transitions. Cell 184, 1914–1928.e19 (2021).

39. Ferme, L. C., Ryan, A. Q., Haase, R., Modes, C. D. & Norden, C. Timely neurogenesis enables increased nuclear packing order during neuronal lamination. 2024.11.12.623216 Preprint at 10.1101/2024.11.12.623216 (2024).

40. Mayran, A. et al. Cadherins modulate the self-organizing potential of gastruloids. 2023.11.22.568291 Preprint at 10.1101/2023.11.22.568291 (2023).

41. Durdu, S. et al. Luminal signalling links cell communication to tissue architecture during organogenesis. Nature 515, 120–124 (2014).

42. Shyer, A. E. et al. Emergent cellular self-organization and mechanosensation initiate follicle pattern in the avian skin. Science 357, 811–815 (2017).

43. Oyler-Yaniv, A. et al. A Tunable Diffusion-Consumption Mechanism of Cytokine Propagation Enables Plasticity in Cell-to-Cell Communication in the Immune System. Immunity 46, 609–620 (2017).

44. Phng, L.-K. et al. Nrarp Coordinates Endothelial Notch and Wnt Signaling to Control Vessel Density in Angiogenesis. Dev. Cell 16, 70–82 (2009).

45. Westerfield, M. The Zebrafish Book: A Guide for the Laboratory Use of Zebrafish (Danio Rerio). (M. Westerfield, 1995).

46. Nüsslein-Volhard, C. & Dahm, R. Zebrafish: A Practical Approach. (Oxford University Press, 2002).

47. Peri, F. & Nüsslein-Volhard, C. Live Imaging of Neuronal Degradation by Microglia Reveals a Role for v0-ATPase a1 in Phagosomal Fusion In Vivo. Cell 133, 916–927 (2008).

48. Marras, S. A. E., Bushkin, Y. & Tyagi, S. High-fidelity amplified FISH for the detection and allelic discrimination of single mRNA molecules. Proc. Natl. Acad. Sci. 116, 13921–13926 (2019).

49. Untergasser, A. et al. Primer3—new capabilities and interfaces. Nucleic Acids Res. 40, e115 (2012).

50. Schindelin, J. et al. Fiji: an open-source platform for biological-image analysis. Nat. Methods 9, 676–682 (2012).

51. Rycroft, C. H. VORO++: A three-dimensional Voronoi cell library in C++. Chaos Interdiscip. J. Nonlinear Sci. 19, 041111 (2009).

52. Lun, A. T. L., McCarthy, D. J. & Marioni, J. C. A step-by-step workflow for low-level analysis of single-cell RNA-seq data with Bioconductor. Preprint at 10.12688/f1000research.9501.2 (2016).

53. Stuart, T. et al. Comprehensive Integration of Single-Cell Data. Cell 177, 1888–1902.e21 (2019).

54. Soneson, C. & Robinson, M. D. Bias, robustness and scalability in single-cell differential expression analysis. Nat. Methods 15, 255–261 (2018).

55. Kobak, D. & Berens, P. The art of using t-SNE for single-cell transcriptomics. Nat. Commun. 10, 5416 (2019).

