## Supplementary Figures, Tables, Notes for "In toto analysis of multicellular arrangement reduces embryonic tissue diversity to two archetypes that require specific cadherin expression"

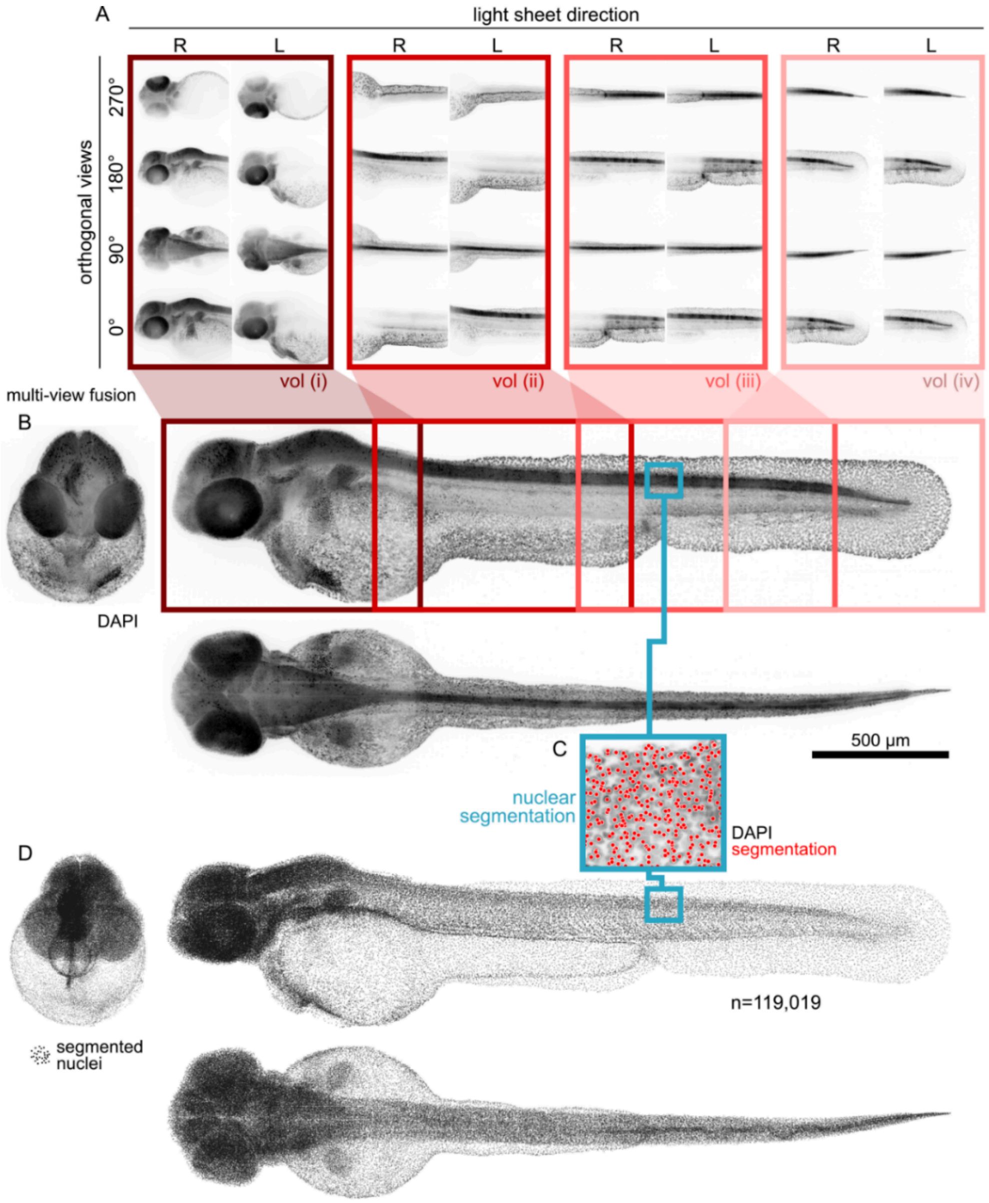


*Supplementary Figure 1: Illustration of multi-view light sheet microscopy and nuclear segmentation pipeline*

***A**: Raw images of zebrafish embryos at 48 hpf with DAPI-labelled nuclei were acquired using multi-view light sheet microscopy. 32 different volumetric image stacks were acquired per sample. Those consisted of four different, overlapping imaging volumes (vol (i) – vol (iv)) with four orthogonal views each (0° – 270°) and two opposing light sheet directions (R, L). **B**: Stacks were iteratively fused using Contrast Limited Adaptive Histogram Equalization* (Hummel, 1977) *and Lucy-Richardson deconvolution* (Lucy, 1974; Richardson, 1972)*, generating one volumetric image of the whole embryo consisting of the highest contrast regions of individual stacks* (Albert, 2021). ***C**: Individual nuclei were localised across the whole fused image using an iterative, multi-level watershed transformation and Gaussian Mixture Models* (Amat et al., 2015; McDole et al., 2018). *Shallow maximum intensity projection across 50 µm. **D**: Point cloud representation of the sample. Locations of the segmented nuclei rendered as points. Images in A, B, D are whole-stack maximum intensity projections.*

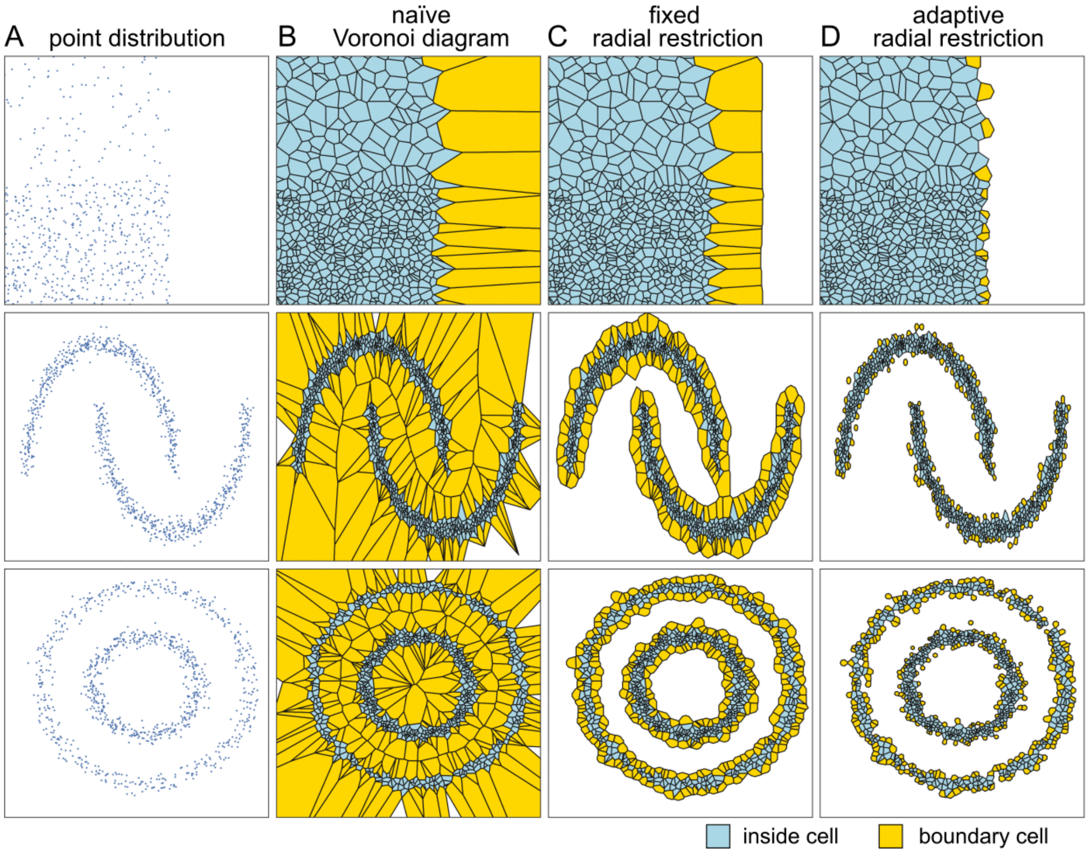


***Supplementary Figure 2: Illustration of adaptively restricted Voronoi diagram***

***A**: Distribution of points that generate the Voronoi diagrams. **B**: Naïve, unrestricted Voronoi diagram. **C**: Voronoi diagram with boundary cells restricted by a maximum radius. **D**: Adaptively restricted Voronoi diagram. In all datasets the naïve Voronoi diagram generates infinitely large boundary cells on the outside and connects unconnected regions on the inside. **Top**: Uniformly random distributed points of low (top) and high (bottom) density. Fixed radially restricted boundary cells are too large, and extend homogeneously to the outside, creating the impression of a smooth surface. Adaptively restricted boundary cells are larger in sparse and smaller in dense regions continuing the shape and size of their neighbouring cells. **Middle**: Double-crescent shaped random point distributions. Radially restricted boundary cells still connect between the crescents and are biased to similar size. Adaptively restricted boundary cells do not connect the crescents and generate isolated cells for points that are too distant from the main objects. **Bottom**: Concentric circle shaped random point distributions. Radially restricted boundary cells are of similar size, while adaptively restricted boundary cells are smaller in the denser inner ring. Note that the shown illustrations were calculated in 2D, while all analysis in this work was performed in 3D.*

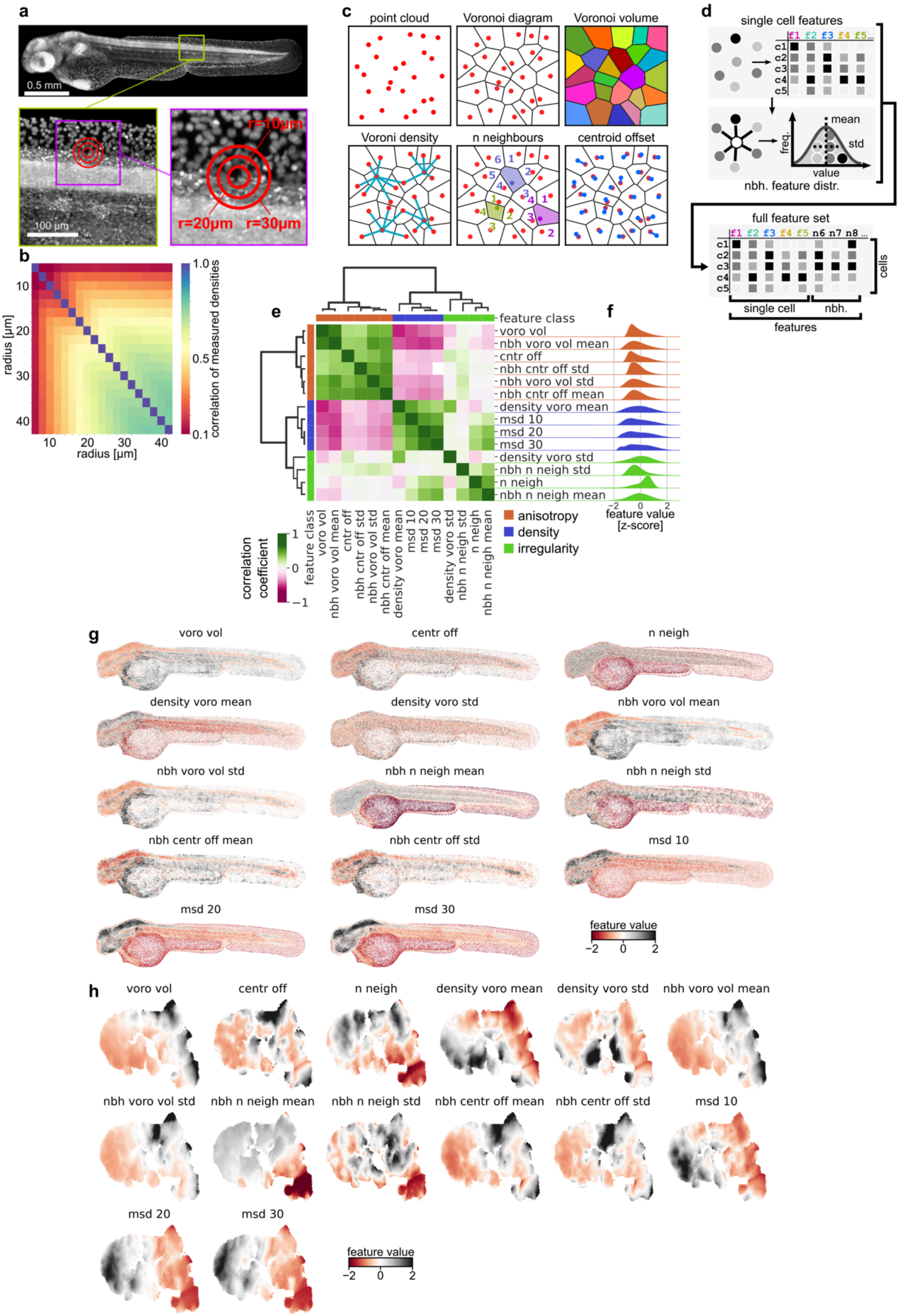


***Supplementary Figure 3: Organisational feature definition, clustering and distributions***

***a**: Illustration of multi-scale kernel density estimation. Density is measured at three different length scales (10, 20, and 30 µm) for every nucleus individually. The density of the lowest radius is measured by counting the number of nuclei inside a sphere of radius 10 µm and dividing this number by the volume of the sphere. For the two larger* *radii, a spherical shell with inner radius 10/20 µm and outer radius 20/30 µm was used; the volume of each corresponding spherical shell was used for normalisation.* ***b****: Correlation of density measurements between different radii. Used data from N=34 samples and n=4,080,605 cells. Selected radii exhibit a correlation of <50%. Lowest radius was selected to avoid bias towards 0 counts and corresponding 0 µm*$^{-3}$ *density measurements.* ***c****: Illustrations of organisational features derived form Voronoi cells. A restricted Voronoi diagram is generated from a point cloud. For each individual point in the cloud the volume of the Voronoi cell, its average inverse distance to all neighbours, the number of neighbours and the distance between seed point and centroid are measured. Note that a three-dimensional Voronoi diagram is used for all measurements.* ***d****: Variability of features are evaluated in local neighbourhood. For Voronoi volume, n neighbours and centroid offset, the mean and standard deviation (std) in the local Voronoi neighbourhood of individual points are evaluated. Together with single cell features (see above) these features are used to characterise the organisation of individual cells in an explicit way.* ***e****: Hierarchical clustering of organisational features based on their autocorrelation indicates three classes. Hierarchical agglomerative clustering was performed on z-scored feature values using Ward linkage* (Ward, 1963) *and Euclidean distances. Note that density and anisotropy features exhibit anticorrelation, indicating some degree of co-linearity. Voro: Voronoi, vol: Volume, nbh: neighbourhood, cntr off: centroid offset, msd: multi-scale density, neigh: neighbours.* ***f****: Global histograms of individual organisational feature values are contained within ±2 standard deviations around the mean. The histograms of density and anisotropy features are dissimilar, indicating that, while they exhibit co-linearity in general, they carry independent information for individual cells. Feature values are z-scored per feature. Colours correspond to the identified feature classes. Histogram density was estimated using a Gaussian kernel estimation with the bandwidth selected via Scott's rule* (Scott, 1992) *multiplied by factor 1.4 to suppress artifacts. N=34, n= 4,080,605.* ***g****: Different features highlight different anatomical regions of the embryo. Renderings of segmented nuclei as dots; z-projected. Colour indicates z-scored feature values. Note that features of one class (e.g. density) exhibit similar, but not identical spatial distributions, indicating that every feature carries unique information.* ***h****: Distribution of individual fieatures on a t-SNE embedding of the organisational feature space. While the distributions of individual features are unique, they are similar between features of a class; intra-class variation is more local. Hexagonal bin plots with 100x100 bins. N=34, n=408,060. Values are mean per bin. Bins with number of cells* $n_{bin} < \frac{n}{(100x100)\cdot 0.3}$ *are not shown to avoid edge effects. voro: Voronoi, vol: Volume, nbh: neighbourhood, cntr off: centroid offset, msd: multi-scale density, neigh: neighbours.*

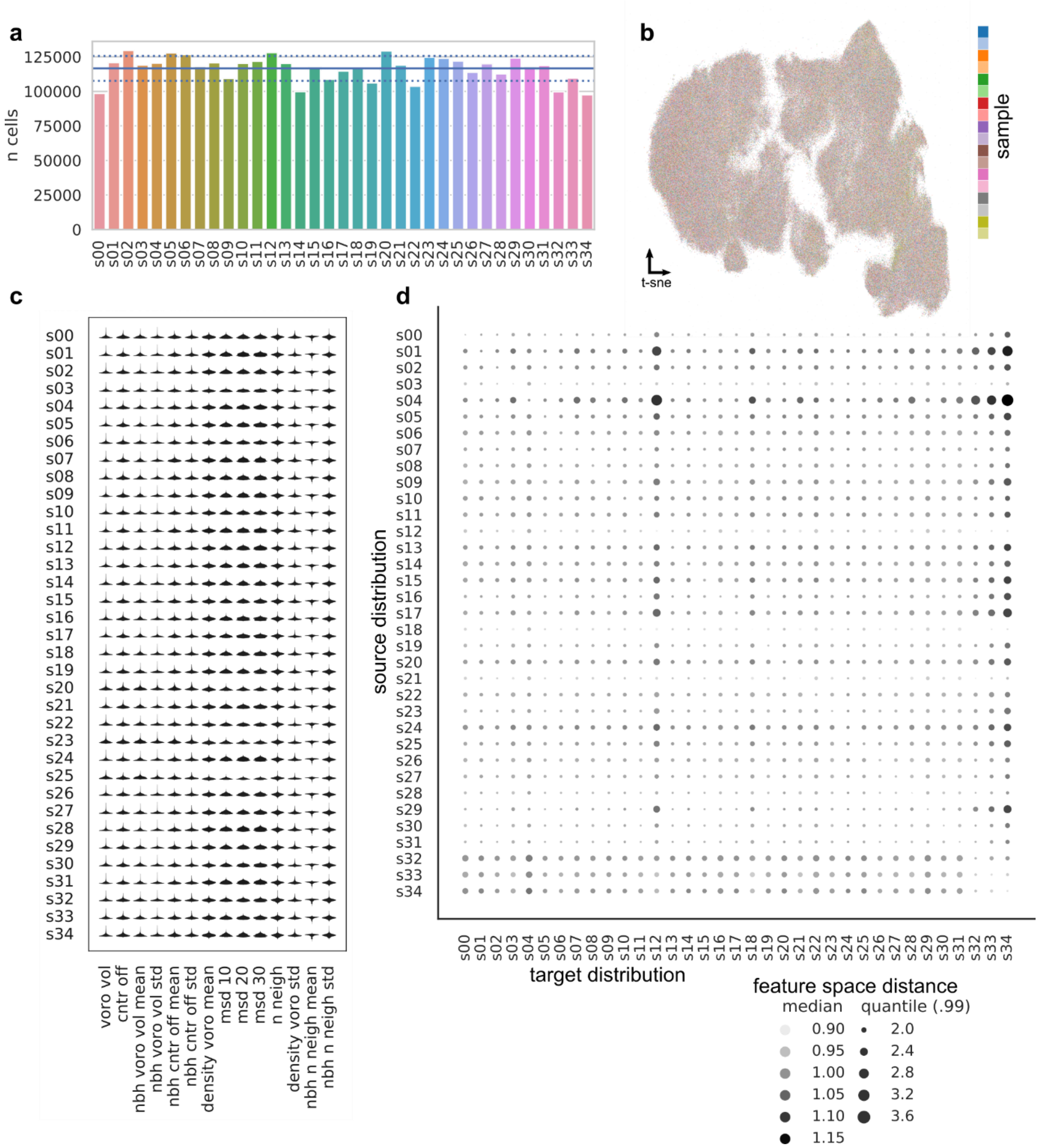


***Supplementary Figure 4: Feature distributions of samples are aligned***

***a**: Number of nuclei are similar between samples. The mean number of cells per sample at 48 hpf was 116,600 ± 9,000 (7.72%); solid lean denotes mean, dashed line mean ± standard deviation. **b**: Organisational feature space is well mixed. Individual samples are coloured individually, however, no region of the t-SNE embedding stands out with a single colour, indicating that, overall, the used samples are similar in terms of their organisational feature distributions. **c**: Distributions of organisational feature values are qualitatively similar between samples. **d**: Distance between samples in organisational feature space. Shown are the distance between single points in the source distribution (y-axis) and their nearest neighbour in the target distribution (x-axis) as total median and the mean of the 1% most distant points. Distance metric: Euclidean distance in 14-dimensional space. Individual samples are labelled sXX (e.g. s01).*

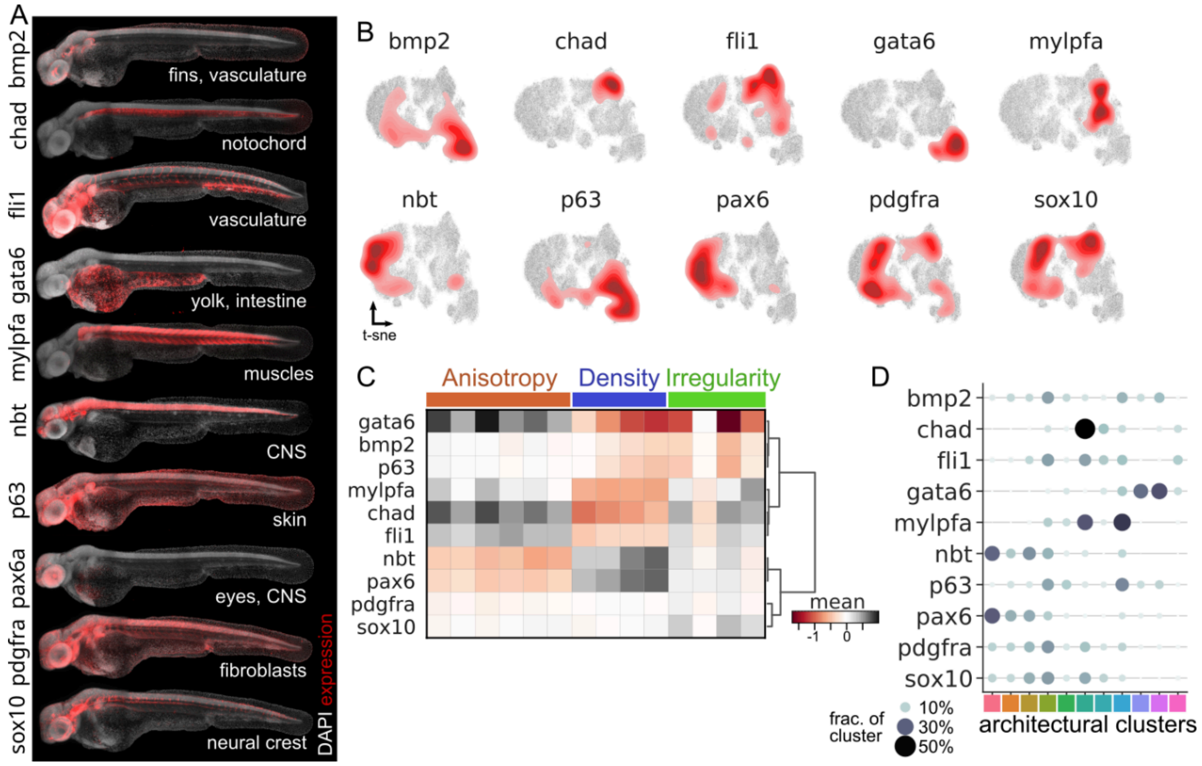


***Supplementary Figure 5: Tissue/cell type specific markers exhibit distinct tissue organisational profiles.***

*A: Spatial distributions of the labelled cells. Maximum intensity projections of in toto multi-view fused light sheet images of 48 hpf zebrafish embryos. Most genes were labelled via HCR RNA FISH, with the exception of nbt and fli1 for which the transgenic fishlines Tg(NBT:dsRed) and Tg(fli1:GFP) were used. Markers were pairwise multiplexed, e.g., chad and mylpfa were stained in the same sample and were imaged in different fluorescent channels. For each marker three replicates were analysed. CNS: central nervous system. B: Mapping of marker positive cells onto a t-SNE of the organisational feature space. Red regions represent the density of cadherin positive cells; grey points are non-expressing and are shown for reference. Density of positive points on embeddings was estimated using a Gaussian kernel estimation with the bandwidth selected via Scott's rule (Scott, 1992). C: Mean organisational feature profiles of marker positive domains. Clustering based on feature similarity links related tissues such as nbt and pax6. Hierarchical agglomerative clustering was performed using Ward linkage (Ward, 1963) and Euclidean distance on z-scored feature values. D: Marker positive domains are composed of unique distributions of organisational clusters. Note that the distribution of architectural clusters adds to 100% for each marker independently of its total abundance and that the distributions do not add to 100% for each cluster.*

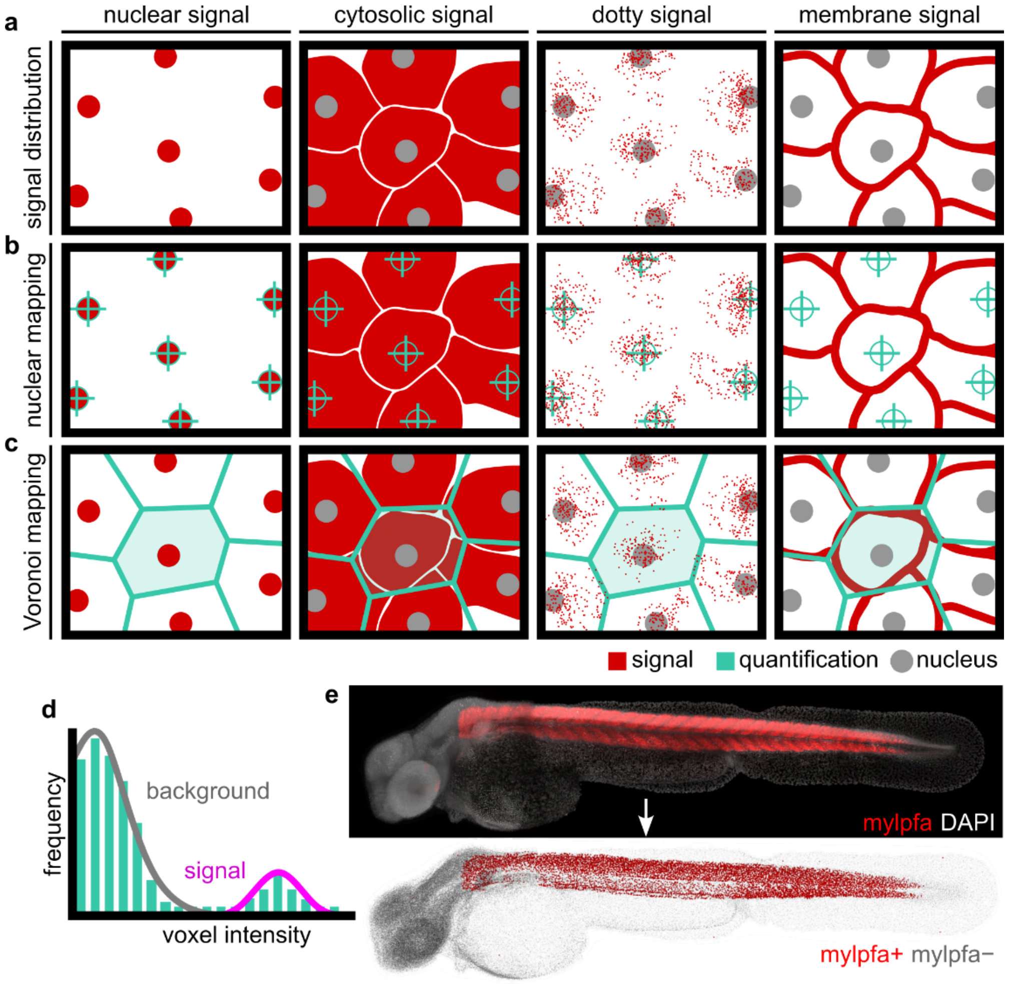


***Supplementary Figure 6: Illustration of intensity classification***

***a**: Different localisations of signals in cells encountered in this study. Depending on the labelling technique or the used transgenic fishlines, localisation of the fluorescent signal within the cell was often externally determined. **b**: Nuclear mapping uses the location of the segmented nuclei to integrate intensities within a spherical region. The radius of that region is a free parameter and needs to be adjusted to the size of the measured object. **c**: Integration within individual Voronoi cells enabled intensity measurements in a larger area in proximity to individual cells. Subsequently, individual pixels were analysed to determine whether a cell was marker positive. **d**: Illustration of the histogram-based pixel classification for the Voronoi integration scheme via Gaussian noise model. Two Gaussian distributions were fitted to the histogram. One captured the noise and one the signal, enabling subsequent analyses to classify cells as positive/negative via global thresholds. **e**: Transformation of an in toto light sheet image of a 48 hpf zebrafish embryo labelled with DAPI and HCR RNA FISH (spot-like pattern) against the muscle specific gene mylpfa into a point cloud representation with mylpfa positive cells highlighted. Maximum intensity projections.*

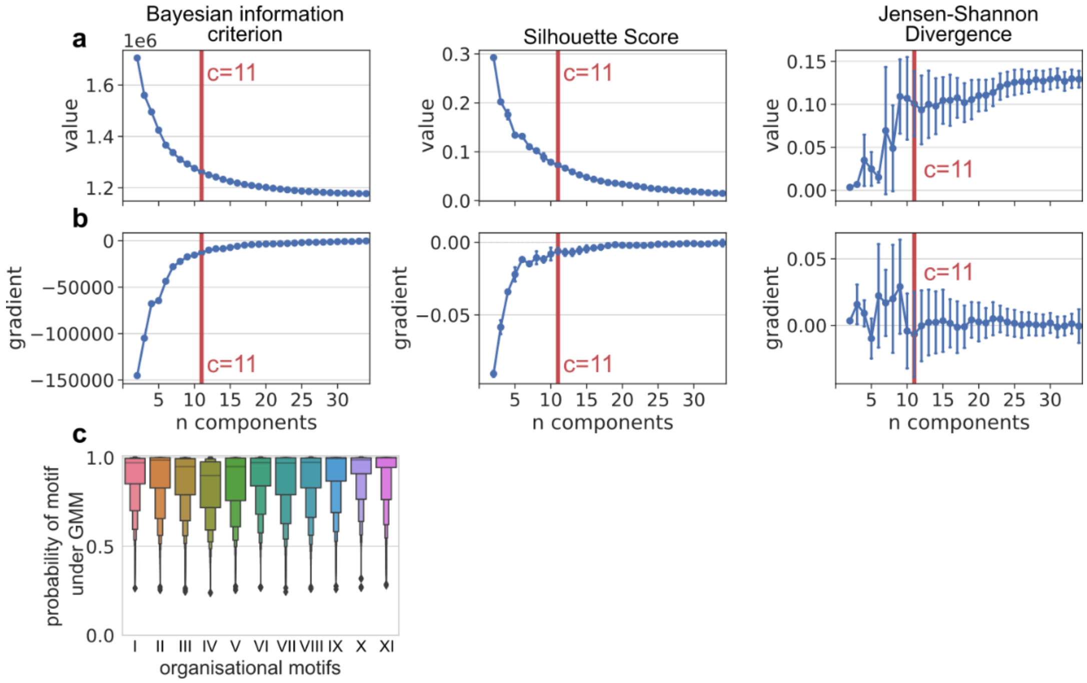


***Supplementary Figure 7: Determination of number of organisational motifs***

*Organisational motifs are determined via Gaussian mixture model (GMM) based on individual cell's organisational feature profiles. To determine the appropriate number of GMM components for the organisational data of 48 hpf zebrafish embryos (N=34, n=4,080,605), GMMs with between 2 and 34 components were generated from 1% randomly selected nuclei and the performance of each GMM was evaluated using three metrics: Bayesian information criterion (BIC), Silhouette score (SilS) and Jensen-Shannon Divergence (JSD) (see Suppl. Note 3 for details on the used metrics). This process was repeated 100 times to estimate variability.* ***a*** *shows the performance of the three metrics and* ***b*** *the gradient of these metrics for consecutive number of components. Error bars denote standard deviation of the mean. Using standard criteria, the number of components is determined to 11* (McLachlan and Rathnayake, 2014; Nielsen, 2019). *For this number of components, the average probability of correct component assignment for each point and component was 97% (median) and never lower than 90% (motif IV), indicating that the identified number of components appropriately models the distribution of nuclei in organisational feature space (**c**).*

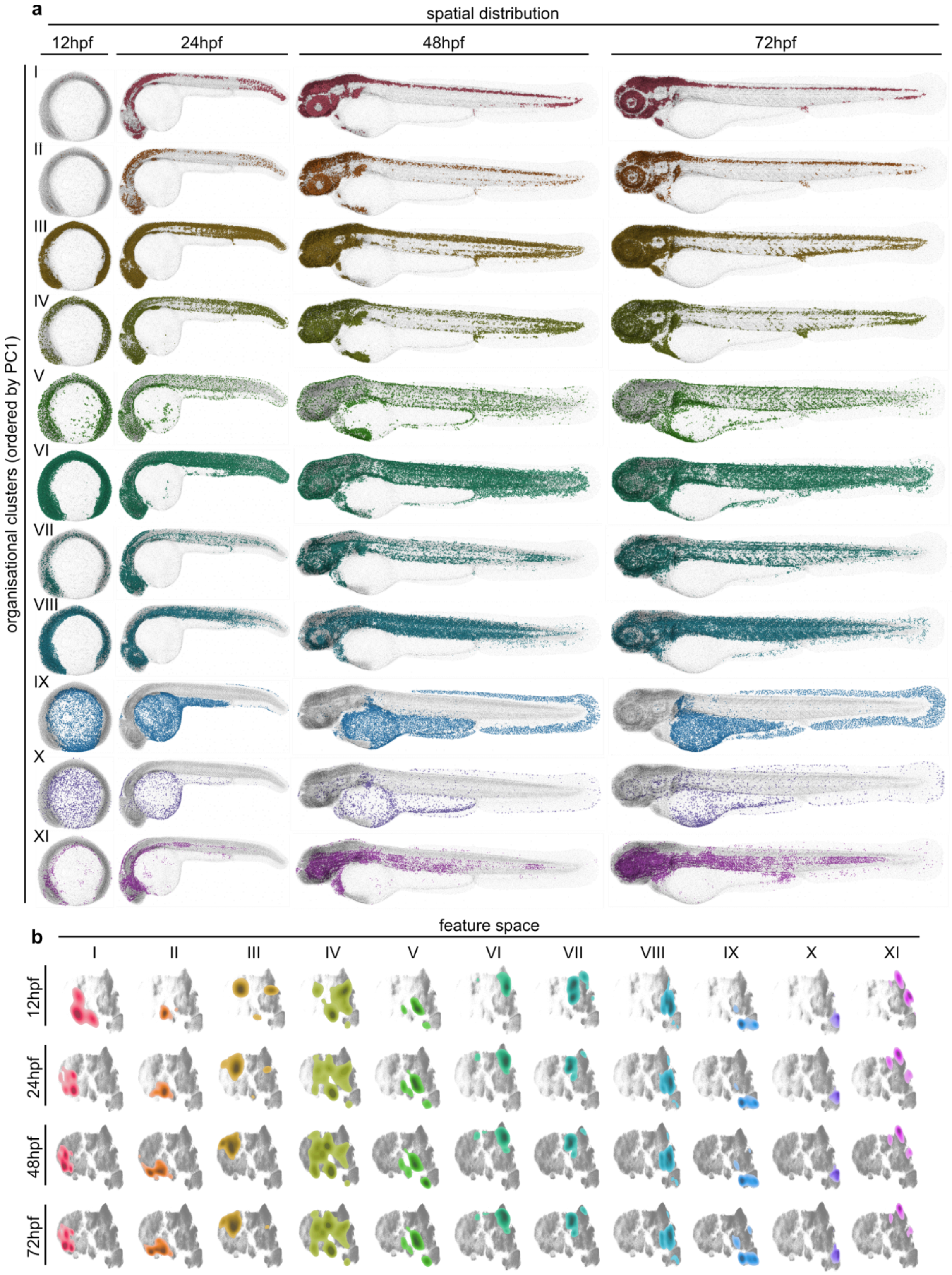


***Supplementary Figure 8: Distribution of organisational motifs***

***a**: Spatial distributions of organisational motifs at different developmental stages. Motifs map to compact, connected regions on the samples that qualitatively encompass different anatomical regions. This highlights the organisational similarity between unrelated tissues. Note that no explicit point coordinates were used during the generation of the motifs. Motifs are ordered by the level of the average first principal component across the whole feature set.* ***b**: Individual clusters at different time points mapped onto a temporally joint feature space. Note that the underlying organisational feature space (grey regions) develops over time, limiting the extents of some motifs. Density of positive points on embeddings was estimated using a Gaussian kernel estimation with the bandwidth* *selected via Scott's rule (Scott, 1992); density is not normalised to total cell number. Grey dots represent cells outside a specific motif. N=16, n=1,698,230*

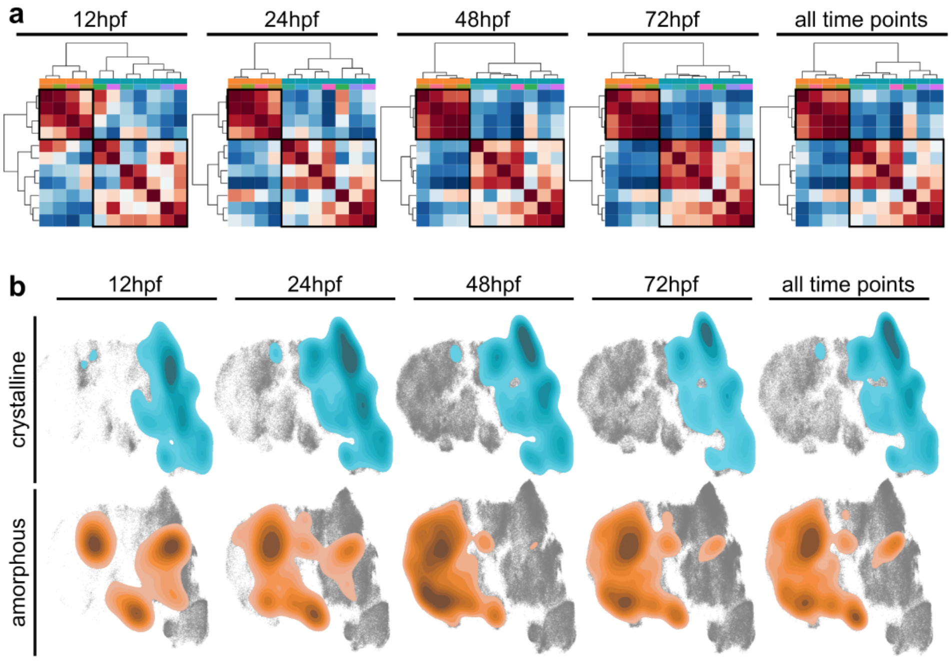


***Supplementary Figure 9: Organisational archetypes exist at all analysed time points***

***a**: Two organisational archetypes are present at every time point individually. Intra-cluster correlation and inter-cluster anticorrelation increase over time. Similarity of organisational motifs based on mean organisational features per motif. Joint analysis of all time points. Hierarchical clustering was performed on z-scored feature values using Ward linkage (Ward, 1963) and Euclidean distances. Colours on the upper two rows indicate archetype and motif association respectively. **b**: The organisational archetypes map to opposite sides of the organisational feature space embedding and show little overlap at every time point individually. Density of positive points on embeddings was estimated using a Gaussian kernel estimation with the bandwidth selected via Scott's rule (Scott, 1992). t-SNE embedding of organisational feature space. N=16 (4 per time point), n=1,698,230*

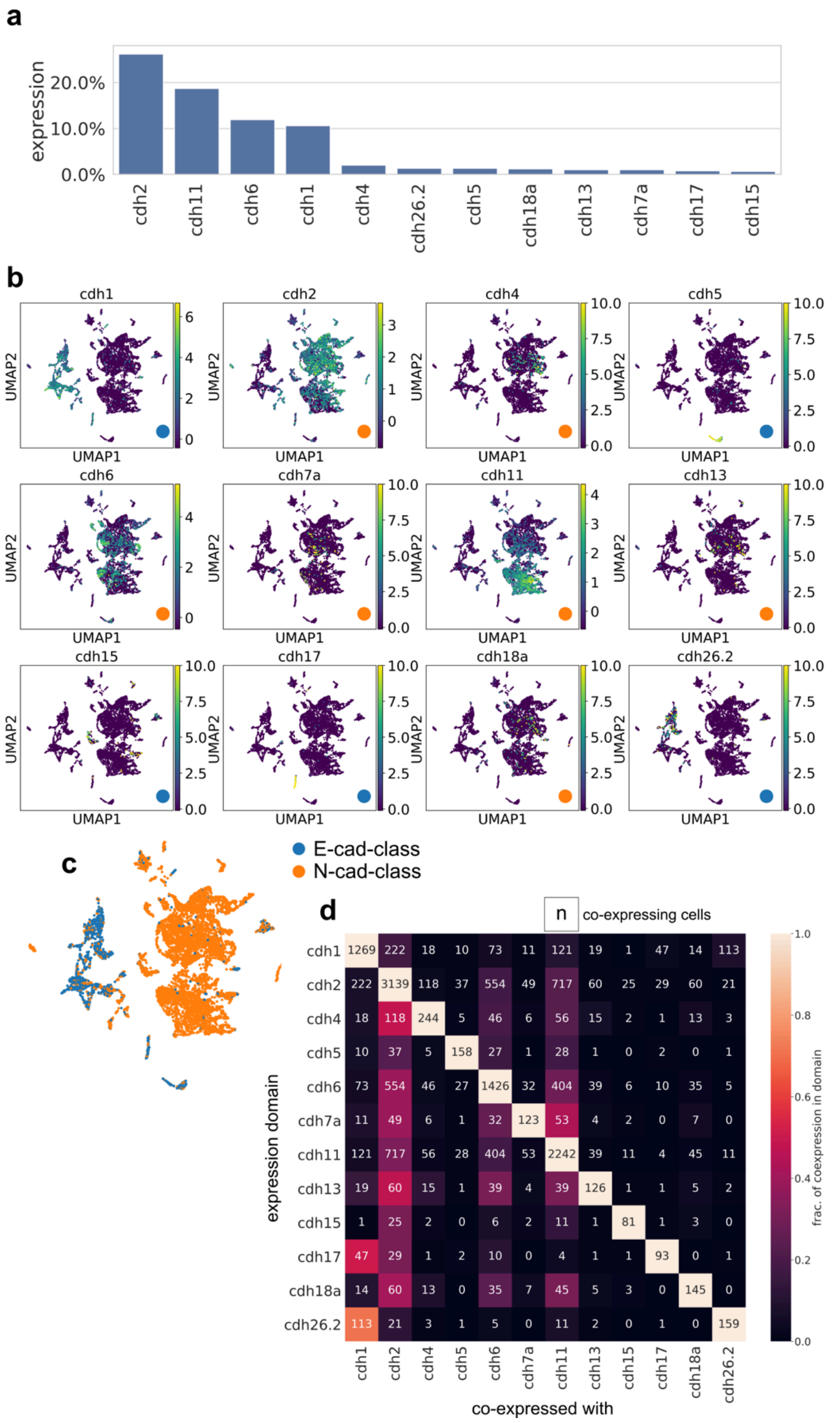


***Supplementary Figure 10: Cadherin expression in scRNAseq analysis exhibits bipartitioning which is not explained by single cell co-expression.***

***a**: Cadherin expressing cells that exceed 1% of all cells in the dataset were selected for analysis. cdh2, cdh11, cdh6, and cdh1 are more broadly expressed than the other domains. Note that individual cells can express multiple cadherins and hence, the total proportion does not add up to 100%. **b**: Individual cadherin domains localise to specific regions of transcriptome embedding. Domains of the E-cadherin-like class (blue) localise to regions that are distinct within the class and between classes, while domains of the N-cadherin-like class co-localise within the class. Scale bars denote the number of transcripts detected per cell. **c**: Localisation of E-cadherin-class and N-cadherin-class cells in transcriptional feature space embedding. Cells E- and*

*N-cadherin-class domains form distinct and predominantly mutually exclusive regions, indicating different transcriptional states. Only cadherin expressing cells are shown. Cells were classified as expressing, if at least one cadherin transcript of was detected. UMAP embedding was calculated on the selected subset using the first 50 principal components of all transcripts according to (Kobak and Berens, 2019).* **d***: Pairwise co-expression of cadherins in scRNAseq dataset. The diagonal entries show the number of cells per cadherin expression domain. Numbers in the rest of the graph indicate the number of cells co-expressing two cadherins. Colouring of the boxes indicates the percentage of cells of one cadherin expression domain (y-axis) co-expressing another cadherin (x-axis). E.g., the orange entry on the bottom left shows that 113 cells co-express cdh26.2 and cdh1; the orange colour reports that 71% of cdh26.2 expressing cells co-express cdh1. The broadest overlap was found between cdh26.2 and cdh1, with 71% of cdh26.2 expressing cells also expressing cdh1. Apart from this overlap, no co-expression exceeded 50% and most cadherins were not co-expressed at significant levels. Importantly, most cdh2 expressing cells do not co-express another cadherin. Data from (Farnsworth et al., 2020), 48 hpf.*

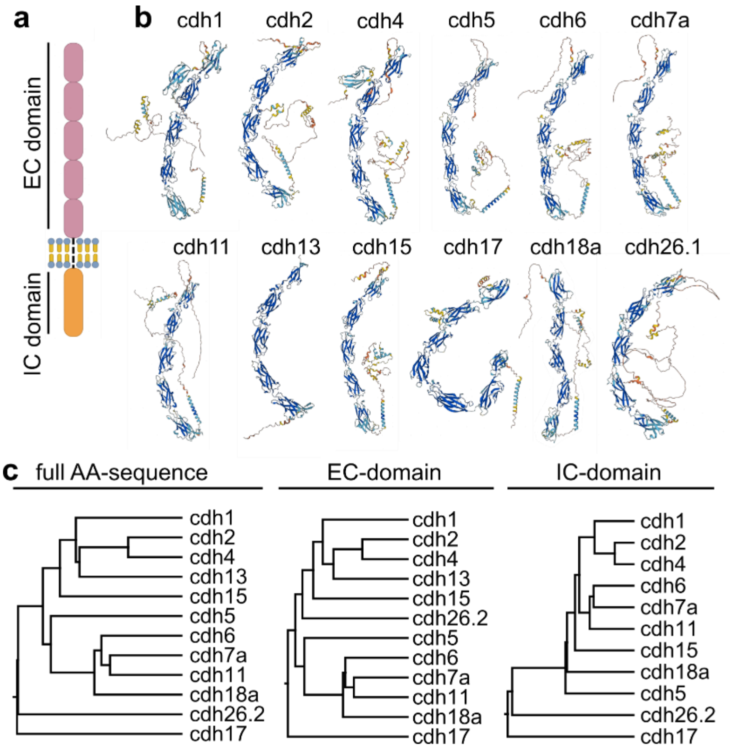


***Supplementary Figure 11: Organisational classification is not predicted by protein similarity***

***a**: Illustration of a generic cadherin protein structure. The extracellular (EC) domain consists of several (commonly five) repeating cadherin domains, while the intracellular (IC) domain has no general stereotypic structure. **b**: Predicted protein structures of the analysed cadherins. Most structures are qualitatively similar and recapitulate the generic model. The most obvious exceptions are cdh1 and cdh17 with six and seven EC repeats. Structures predicted by Alphafold (DB version 2022-11-01) (Jumper et al., 2021; Varadi et al., 2022). **c**: Similarity of cadherin amino acid sequences does not recapitulate clustering based on transcriptional or organisational features. Alignments are based on the cadherin's full-length sequences, their extracellular domains only and their intracellular domains only. Trees built using BLOSUM85 cost matrix, Jukes-Cantor genetic distance model and UPGMA via geneious v2022.0 (Kearse et al., 2012).*

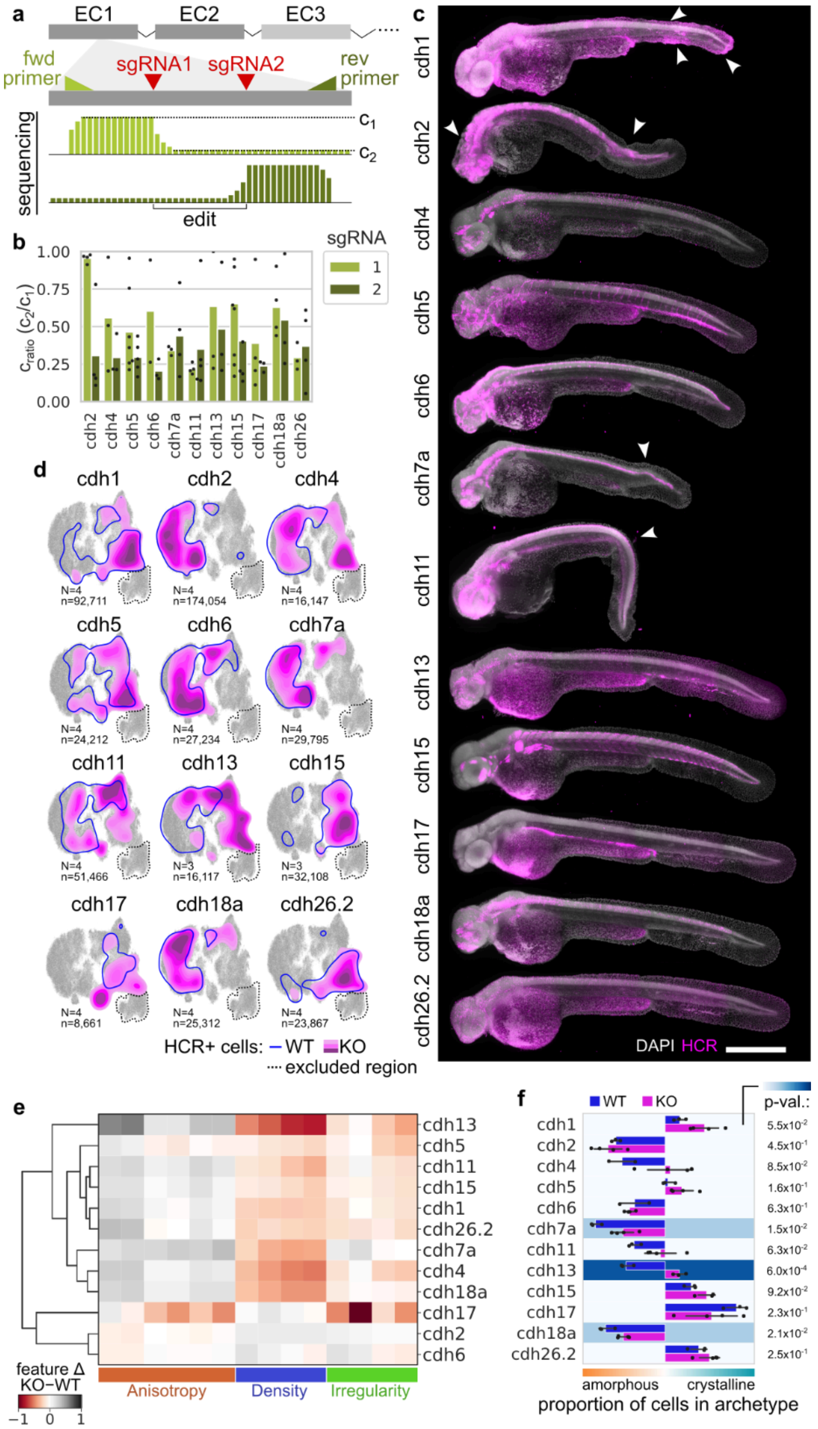


***Supplementary Figure 12: Systematic knockout of individual cadherins using F0 CRISPR/Cas9***

*__a__: Transient 'null mutants' are generated by targeting two sites of the target cadherin's EC1 or EC2 domain with sgRNAs. Successful knockout (frame shift or large deletion) was confirmed via Sanger sequencing across the sgRNAs' homology region. A drop in read quality in the chromatogram following a sgRNA target site indicated a successful edit. $c_1$: chromatogram level pre-sgRNA; $c_2$: chromatogram level post-sgRNA; in direction of the sequencing primer. __b__: Transient knockout of individual cadherins was efficient. $c_{ratio}$ ($c_2$ / $c_1$) measures the drop in sequencing quality following a sgRNA cut site. Values range from 1 (no edit) to 0 (no signal post site). sgRNA1 of cdh2 did not cause a genome edit. All other sgRNAs edited the genome on average. Whole-embryo cdh1 KO was lethal; consequently, verification via sequencing was not possible; cdh1 was knocked out in a mosaic fashion by 8-cell stage injection. __c__: Deletion of individual cadherins causes gross phenotypes for cdh1, cdh2 cdh7a and cdh11. The knockout of the other cadherins did not induce obvious embryo-wide effects. Maximum intensity projections of in toto light sheet images. Nuclei are stained with DAPI and the transcripts of the target cadherins are visualised using HCR RNA FISH. Arrowheads indicate phenotypic regions. __d__: Knockout of cadherin induces* *organisational change in transcript-expressing cells. Distributions of cadherin transcript positive cells on t-SNE representation of organisational feature space. Solid magenta regions are cells from the knockout condition, blue outline shows corresponding regions in wildtype samples (see **Figure 3**), grey regions are non-expressing cells for reference. Regions within the dotted line (cells on yolk) were not included in the analysis due to unspecific labelling of the yolk. **e**: Feature difference between cadherin transcript positive cells in knockout and wildtype. Most domains exhibit a notable shift in organisational features. **d**: Organisational archetype association of cadherin transcript positive cells in wildtype and knockout. Background colour indicates the difference between mean archetype proportions in knockout and wildtype. Most cadherin knockouts organise more crystalline. Error bars indicate 95% confidence interval. Sample number as indicated in D (KO) and **Figure 3** (WT).*

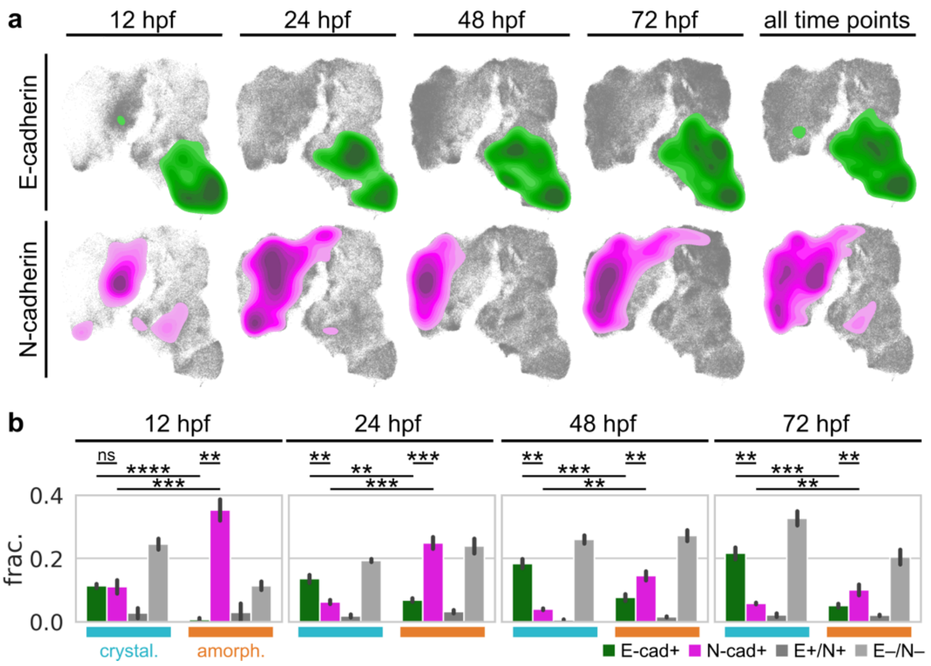


***Supplementary Figure 13: E- and N-cadherin associate with crystalline and amorphous archetype during early development***

***a**: E- and N-cadherin localise on opposing ends of organisational feature space at all analysed time points, indicating consistently different tissue organisation. **b**: E- and N-cadherin consistently associate with crystalline and amorphous archetypes. Fraction of E-/N- cadherin expressing cells that organise crystalline/amorphous is significantly higher than both E-/N-cadherin expressing cells that organise amorphous/crystalline, and N-/E-cadherin expressing cells that organise crystalline/amorphous. Statistical test: Welch's t-test independent samples with Bonferroni correction, p-values: ns: 5.00e-02<p<=1.00e+00, *: 1.00e-02<p<=5.00e-02, **: 1.00e-03<p<=1.00e-02, ***: 1.00e-04<p<=1.00e-03, ****: p<=1.00e-04*

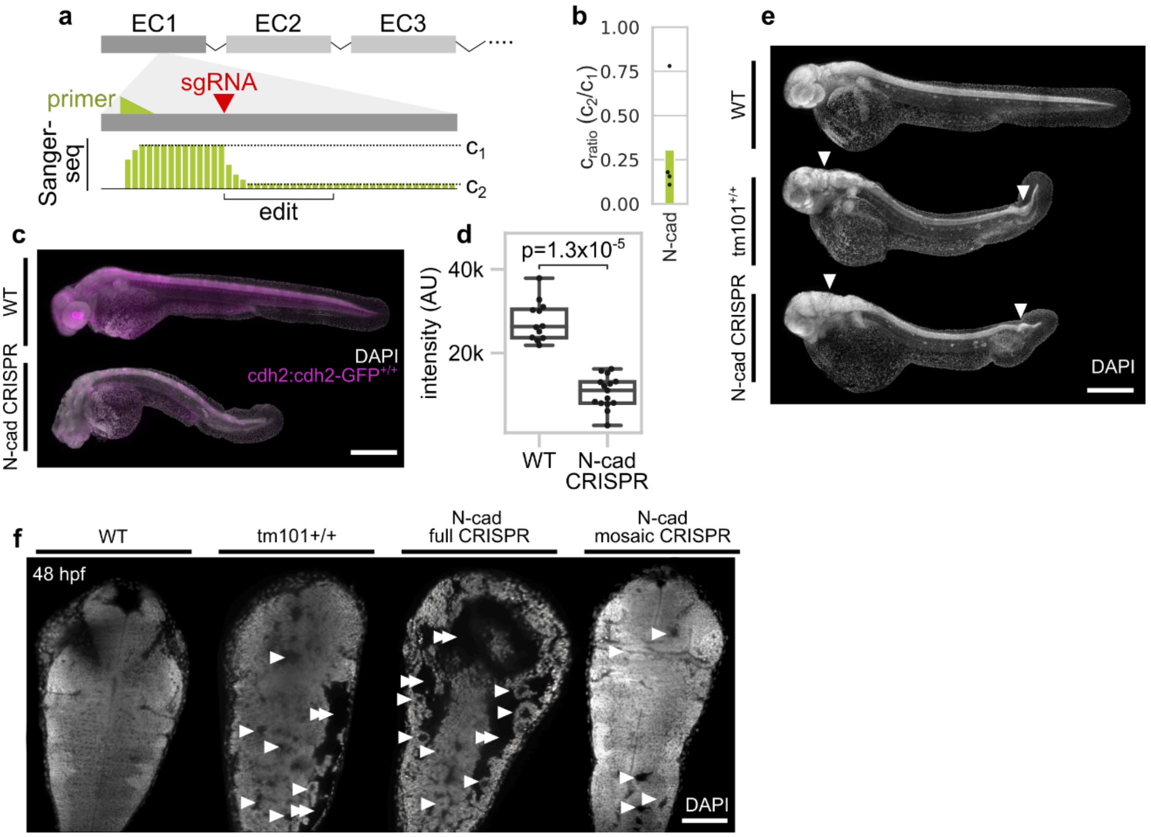


***Supplementary Figure 14: CRISPR mediated, transient KO of N-cadherin is verified by sequencing and imaging, and phenocopies established stable mutant.***

***a**: sgRNA for N-cadherin targets region within the first extracellular repeat (EC) domain. Edit region is amplified via PCR using a primer in proximity of the sgRNA target site and Sanger sequenced. Alignment to the target region exhibits drop in quality (c1 vs. c2) at the target site if edit was successful (**b**), rendering the protein non-sensical. **c, d**: Transient N-cadherin KO (N-cad CRISPR) leads to loss of N-cadherin fusion protein (BAC(cdh2:cdh2-GFP)) expression. Quantification based on n=12 WT samples and n=15 N-cad CRISPR samples. Statistical test: Mann-Whitney. **e**: N-cad CRISPR phenocopies the* parachute *mutant ($cdh2^{tm101}$). Arrowheads indicate regions exhibiting the strongest phenotype in the brain and tail. **f**: KO of N-cad via CRISPR injection into 1-cell stage of embryonic development (full CRISPR) results in severe brain phenotype including loss of brain regions, cyst formation (arrowheads) and dramatic ventricular enlargement (double arrowhead). The parachute mutant (tm101) exhibits a similar, albeit slightly weaker phenotype. KO of N-cad via CRISPR injection at the 8-cell stage (mosaic CRISPR) largely omits this phenotype and leads to only minor defects in brain development compared to WT, tm101+/+ and full CRISPR (arrowheads). Scale bars: c, e: 400 µm, f: 100 µm*

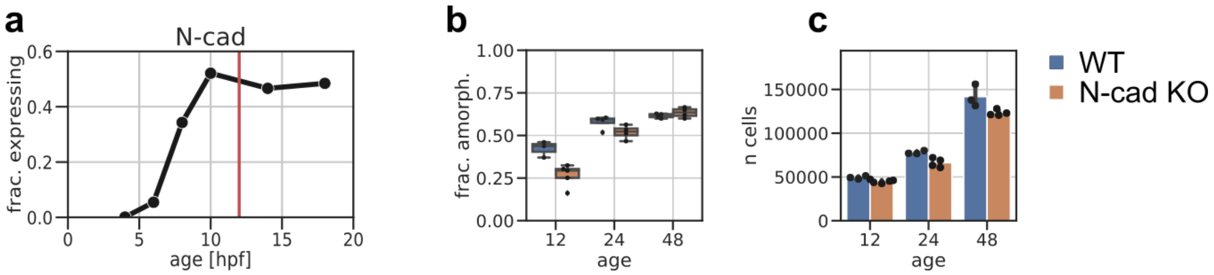


***Supplementary Figure 15: Early N-cad expression and archetypal dynamics.***

***a:*** *N-cadherin expression starts around 6 hpf of zebrafish development and rises sharply until 10 hpf, where it plateaus. Red line demarks 12 hpf, the first time point, where phenotypic analysis of N-cad CRISPR was conducted. Data from (Wagner et al. 2018).* ***b:*** *Fraction of amorphously organising cells between wildtype and CRISPR mediated N-cad knockout is lower at 12 hpf but nears parity at 48 hpf.* ***c:*** *Number of cells per sample is the same between WT and N-cad KO at 12 hpf. KO shows reduced cell count at 24 and 48 hpf, indicating gross developmental defects.*

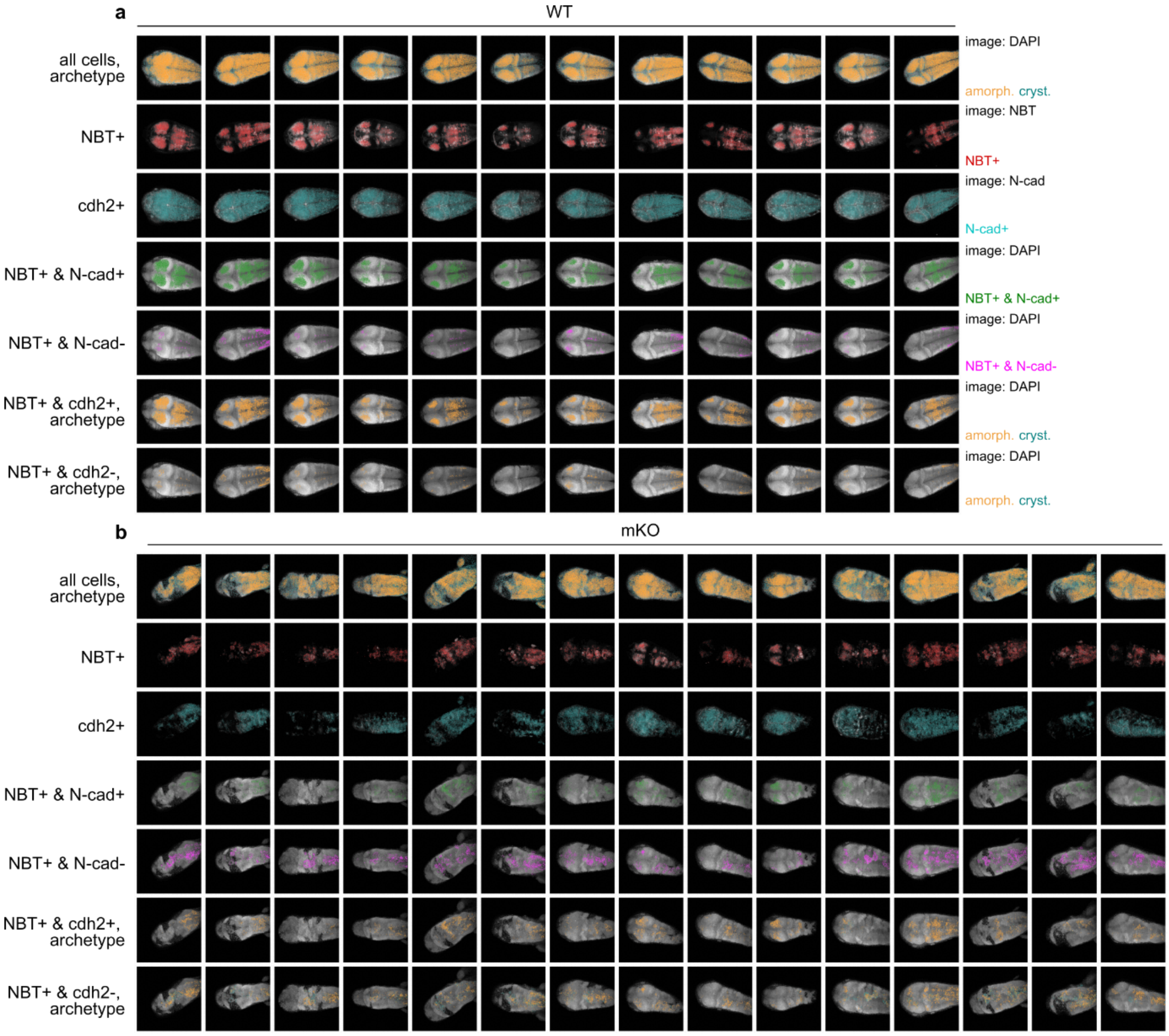


***Supplementary Figure 16: Mosaic CRISPR of N-cadherin generates identifiable knockout and induces morphological phenotype***

*Maximum intensity projections of all imaging data used for the mosaic N-cadherin knockout experiments. Wildtype samples in* ***a*** *exhibit conventional tissue architecture in the brain, express NBT in a regular restricted pattern and express N-cadherin in a wider pattern. Classification of segmented nuclei based on gene expression works reliably. Co-expression domains of NBT and N-cadherin are regular, bilaterally symmetrical, and predominantly organised amorphously. Cells expressing only NBT are almost fully absent. mKO samples in* ***b*** *exhibit a visible morphological phenotype. Cells expressing only NBT are more frequent and organise more crystalline.*

### Supplementary Notes

**Supplementary Note 1: Adaptively restricted Voronoi diagram**

To generate features that captured the spatial organisation of the segmented nuclei, we needed to establish spatial neighbourhood relationships between them in a manner that reflected local cell arrangement. We used a modified version of the three-dimensional Voronoi diagram to achieve that. In a conventional Voronoi diagram, space is partitioned based on the distance between neighbouring points, by assigning all space that is closest to an individual point to its 'Voronoi cell', which consequently reflects local organisation in a robustly quantifiable way. This generates *interfaces* between Voronoi cells, that are equidistant between two points, and *vertices*, that are closest to three or more points. One challenge with the Voronoi approach is that it breaks down at external boundaries, since there Voronoi cells grow infinitely large (Suppl. Fig. 2 a,b).

A naïve approach to circumvent this would be to limit the size of Voronoi cells such that they cannot exceed a certain radius or volume. While this method is commonly used to remove giant Voronoi cells e.g., for visualisation, it is not suited for our purpose, since it biases boundary Voronoi cells to a pre-defined, maximal volume, making them incomparable to Voronoi cells from the inside. In fact, such cells differ significantly even from their close neighbours that are just one layer below (Suppl. Fig. 2 c).

Another approach to limit boundary Voronoi cells is to generate a surface mesh of the whole point cloud and to cut the boundary Voronoi cells with it (Sainlot et al., 2017; Wang et al., 2020; Yan et al., 2010). Given a high-quality surface mesh, this method is very powerful, as it would conserve the surface shape of the sample. However, generating such a mesh from imaging data is difficult and would require specific genetic markers that may not be generally attainable. Deriving a surface mesh from the point cloud itself is also generally challenging. For point clouds with convex topology a surface mesh can be trivially generated from the convex hull. However, concave topologies cannot be isolated in such a rigorous way and would involve the use of parameterised methods such as alpha shapes that require tuning of 'concavity' parameters or similar (Gardiner et al., 2018). Moreover, conventional meshing approaches are only sensitive to boundaries to the outside and would fail to detect cavities inside the point cloud (Abdelkader et al., 2020).

To overcome these limitations, we developed a method in which individual Voronoi cells were adaptively size- and shape restricted using auxiliary seed points estimated from their neighbourhood. Initially, a naïve radially restricted Voronoi diagram was generated with restriction radius 50 µm, and boundary Voronoi cells were defined as cells with at least one face which was not shared with another Voronoi cell. For these boundary cells, all neighbouring cells were identified and their average number of neighbours $n_n$ as well as their average volume $V_n$ was determined. From this, $n_n$ auxiliary points were placed isotopically, in the pattern of a Fibonacci lattice (González, 2009), on a sphere around every boundary nucleus at the distance $d_n = 2 \cdot \sqrt[3]{3V_n/(4\pi)}$. Subsequently, all auxiliary points that were placed outside the boundary Voronoi cell were deleted. This way auxiliary points were only placed in spots where they would restrict the Voronoi diagram towards the boundary. Finally, a second Voronoi diagram was generated on the combined set of nuclei and auxiliary points, resulting in restricted boundary cells (Suppl. Fig. 2 d). The auxiliary points were not considered for further processing.

To increase robustness, a k-nearest neighbour search was implemented in case a cell was surrounded by exclusively boundary cells. In that case, nearest neighbours would be iterated over starting with the closest and the $n_n$ and $V_n$ would be generated from the first five non-boundary nearest neighbours. This method generated a Voronoi diagram in which boundary cells, and cells in sufficiently large cavities were restricted in a way that continued the distribution of the local neighbourhood. This enabled the comparison of inside and boundary cells in a common framework. Beyond the scope of this study, this method may generally be useful to construct restricted Voronoi diagrams on arbitrary point clouds without boundary mesh since it only requires the upper limit of a Voronoi cell's size as a single free parameter.

**Supplementary Note 2: Definition of organisational features**

***Multi-scale cell density***

A fundamental feature of tissue organisation is cell density. We measured cell density at different length scales to not only assess the compactness of a cell's immediate neighbourhood, but also the overall density of its surrounding tissue. To achieve this, we employed a kernel density estimation. For an individual cell, we centred on the coordinates of its nucleus and counted the number of segmented nuclei in a number of spheres with increasing radii (Suppl. Fig. 3 a). Then, we normalised the measured cell count by the analysed volume to make the values comparable between radii and convert the count into a density value. To identify an appropriate set of radii, we evaluated the correlation between measured densities at different radii. This indicated a strong correlation between consecutive spheres. Measuring cell density in stacked spherical shells rather than solid spheres decreased the correlation between measurements at different radii. Therefore, we chose this geometry as the basis of multi-scale density estimation. Moreover, we selected the radii 10 µm, 20 µm and 30 µm. These were selected since cells have a stereotypic diameter of ~10 µm, which led to a bias towards 0 $\mu m^{-3}$ measurements in low-density areas for smaller radii, and spherical shells exceeding a radius of 30 µm exhibited correlation exceeding 60%, indicating mutual information content for larger radii (Suppl. Fig. 3 b).

Summarising, we estimated local cell density per segmented nucleus by counting the number of segmented nuclei within spherical shells between 0-10 µm (solid sphere), 10-20 µm, and 20-30 µm, normalising the count by the measured volume. This yielded a local, volumetric density estimation $[\,]_{msd}$ following the equation

$$[\,]_{msd} = \frac{3\, n_{points}}{4\,\pi(R^3 - r^3)}$$

with the number of counted points per volume points $n_{points}$, the inner sphere radius $r$ and the outer sphere radius $R$. Evaluating cell density at different length scales was key to evaluate, for example, whether densely packed cells were a compact cluster in an otherwise sparse context, or if the dense packing was the overall tissue organisation.

***Voronoi cell features***

We used the restricted Voronoi diagram to quantify the local distribution of segmented nuclei. The Voronoi diagram is specifically appropriate for this, as it encodes the local distribution of points in the shape and size of individual Voronoi cells. Therefore, quantifying those parameters effectively characterises the local point distribution in a point-by-point way. Moreover, the Voronoi diagram generates a spatial hierarchy, i.e., relationships between points, in a mathematically rigorous way, making it both robust and generally applicable. The following organisational features were measured directly on Voronoi cells and were together used to characterise the local organisation of individual cells.

*Voronoi cell volume*

The volume of a Voronoi cell anti-correlates with the density of its neighbourhood, with the volume being lower in denser regions and higher in less dense regions. Moreover, it also measures the anisotropy of the local neighbourhood since it will be lower if density is high in only one direction. One way to illustrate this would be to imagine a Voronoi cell as an ellipsoid whose volume $V_{ellipsoid}$ would be given by $V_{ellipsoid} = \frac{4}{3}\pi abc$ with $a,\ b,\ c$ being the length of the three semi-axes. If any one of them gets close to 0, the whole volume will go to 0.

The volume of the Voronoi cells was calculated via a tetrahedral decomposition (Bukenberger and Lensch, 2021) of the Voronoi cell. The volume of the tetrahedra was subsequently determined via

$$V_{tetra} = \frac{|\tilde{a} \cdot (\tilde{b} \times \tilde{c})|}{6}; \qquad \tilde{a},\ \tilde{b},\ \tilde{c} = [a,\ b, c] - d$$

with $a,\ b,\ c, d$ being the vectors pointing to each vertex of the tetrahedron. The sum of all tetrahedral volumes was the total Voronoi cell volume (Suppl. Fig. 3 c).

*Voronoi density*

To estimate local cell density in a non-kernel-based way, we measured the distance between individual nuclei and their adjacent nuclei, i.e., nuclei of Voronoi cells sharing an interface (Suppl. Fig. 3 c). Sharing an interface in the Voronoi diagram is equivalent to being connected on the Delaunay graph. The average inverse distance between cells in a Delaunay neighbourhood described local cell density, while the standard deviation of that distribution measured the regularity of density, with higher values indicating more irregularity.

*Number of neighbours*

Since the Voronoi diagram establishes neighbourhood relationships between unstructured points, the number of neighbours of any given point is a key feature. We measured the number of neighbours by counting the number of connections of a point (node) on the Delaunay graph, i.e., the number of shared faces per Voronoi cell (Suppl. Fig. 3 c). Note that faces to the outside, i.e., restricted faces, were not counted. Shared interfaces were not weighted by interface area and, therefore, it was more likely for an irregularly arranged neighbourhood to exhibit higher number of neighbours.

*Centroid offset*

To measure the isotropy of local point arrangements, we measured the Euclidean distance between a Voronoi cell's seed point—the location of the nucleus—and its centroid. The location of the centroid was determined via tetrahedral decomposition of the Voronoi cell. In an isotropic environment, the seed point and centroid of a Voronoi cell are identically localised, while their distance increases as function of local anisotropy (Suppl. Fig. 3 c).

***Quantification of local organisational variability***

To evaluate the variability of feature values in a local neighbourhood, we analysed the distribution of Voronoi cell volume, number of neighbours, and centroid offset in the Voronoi neighbourhood of each segmented nucleus. From that, we calculated mean and standard deviation of the distributions as additional feature values. Importantly, the segmented nucleus used to generate that neighbourhood was excluded from the analysis. This way we were able to contrast the feature value of an individual cell, e.g., its number of neighbours, with the average feature value of the neighbourhood. Moreover, we gained access to the overall feature variability in a cell's neighbourhood via the standard deviation (Suppl. Fig. 3 d). Overall, this neighbourhood analysis expanded our feature set by six neighbourhood features that mainly carry information about local heterogeneity.

***Normalisation enables cross-feature comparison***

To enable further analysis, such as low dimensional embedding and clustering, and to make features comparable between each other, individual feature values were normalised using the standard z-scaling approach:

$$\widetilde{f}_i = \frac{f_i - \langle f \rangle}{\sigma_f}$$

With the individual feature value $f_i$, the normalised feature value $\widetilde{f}_i$, the mean feature value across the samples $\langle f \rangle$ and the corresponding standard deviation $\sigma_f$. We found this scaling approach more robust than a scaling of the minimum and maximum feature value between 0 and 1 since it is less biased towards outliers. The feature normalisation was performed jointly on all samples to conserve sample-to-sample variability and differences between experimental conditions.

**Supplementary Note 3: Gaussian Mixture Model stratifies organisational feature space into eleven motifs**

To identify organisational motifs, we clustered cells based on the similarity of their organisational features. A Gaussian Mixture Model (GMM) was used to approximate the distribution of cells in the organisational feature space. In short, such a model uses a pre-defined number of n-dimensional Gaussian distributions and optimises their coordinates as well as their covariance matrices to best fit a given point distribution. In this context, points are treated as observations, while the GMM models the underlying probability density function. The parameters of the GMM are optimised to maximise the likelihood of the observations under the probability density function. (Reynolds, 2009)

We elected to use a GMM for several reasons. Compared with simpler methods such as K-Means clustering, GMMs generally identifies clusters in data more reliably and is able to model finer-grained distributions more accurately (Patel and Kushwaha, 2020). Moreover, the probabilistic nature of the GMM enabled us to quantitatively assess the model performance on our data. More advanced clustering techniques such as DBSCAN (Ester et al., 1996) or OPTICS (Ankerst et al., 1999) are most sensitive to discrete clusters and hence were unable to identify meaningful clusters in our largely continuous data. Moreover, those methods were computationally significantly more expensive than the GMM.

The free parameter of a GMM is its number of components, i.e., how many Gaussian distributions will be used to approximate a given point distribution. Setting this number correctly is crucial for a good model fit. To determine the number of components, we optimised GMMs with number of components between 2 and 34 and measured the model performance at every iteration by analysing the model's Bayesian Information Criterion, its Silhouette Score and its Jensen-Shannon divergence (Lavorini, 2018).

The Bayesian Information Criterion BIC was defined as

$$BIC = kln(n) - 2ln(\hat{L}(x))$$

with the maximised likelihood function $\hat{L}$, observations $x$, number of model parameters $k$ and the number of observations $n$ (Wit et al., 2012). Low BIC scores indicate a good model fit. However, reportedly, the penalty term of the BIC score (left) is insufficient to suppress a bias towards models with too many components (Lavorini, 2018). Therefore, we evaluated the gradient of the BIC between consecutive numbers of components.

The Silhouette Score was defined as the mean Silhouette Coefficient $s_i$ across all points in each cluster.

$$s_i = \frac{b_i - a_i}{max(a_i, b_i)}$$

With the mean intra-cluster distance of the point $a_i$ and the nearest-cluster distance $b_i$ (Rousseeuw, 1987). Consequently, the Silhouette Score is in the range [-1, 1] with values close to 1 indicating a good fit of the clustering model. Similarly to the BIC, the Silhouette Score often biases analysis towards complex models, which can be circumvented by analysing its gradient.

For the Jensen-Shannon divergence, the dataset was randomly split into two sets of equal size and an n-component GMM was generated on each of them individually. The probability distributions $P$, $Q$ of the two GMMs were estimated by converse random sampling of 20,000 points, i.e. for set $X$ drawn from $P$, the log probabilities were $\log log\,(P(x))$ and $\log log\,(Q(x))$ with $x \in X$, and for set $Y$ drawn from $Q$, the log probabilities were $\log log\,(P(y))$ and $\log log\,(Q(y))$ with y$\in Y$. From this the Jensen-Shannon divergence was calculated as

$$JS = \frac{1}{2} \cdot \left( (2) + \langle \log log\,(P(x)) \rangle_{x \in X} - \langle \log log\,(P(x) + Q(x) \rangle_{x \in X} + \langle \log log\,(Q(y)) \rangle_{y \in Y} - \langle \log log\,(P(y) + Q(y) \rangle_{y \in Y} \right)$$

With the mean over a set $\langle\,\rangle_{x,y \in X,Y}$. A low $JS$ indicates that the two GMMs of the split dataset are similar and hence the number of components fits well with the data (Nielsen, 2019).
To reliable estimate the number of components, the consensus of these three metrics over 100 iterations based on 80,000 randomly selected datapoints was used. From this analysis we determined the number of GMM components to be eleven (Suppl. Fig. 7).

### Supplementary Tables

***Table 1: Parameter values used for TGMM 2.0 nuclear segmentation***

*The parameters ‘imageFilePattern’ and ‘backgroundThreshold’ were set for each preprocessed image individually.*

| Parameter | Value |
|---:|---:|
| anisotropyZ | 1 |
| persistanceSegmentationTau | 5 |
| betaPercentageOfN_k | 0.05 |
| nuPercentageOfN_k | 1 |
| alphaPercentage | 0.7 |
| maxIterEM | 100 |
| tolLikelihood | 1.00E-06 |
| regularizePrecisionMatrixConstants_lambdaMin | 0.02 |
| regularizePrecisionMatrixConstants_lambdaMax | 0.1 |
| regularizePrecisionMatrixConstants_maxExcentricity | 9 |
| temporalWindowForLogicalRules | 5 |
| thrBackgroundDetectorHigh | 1.1 |
| thrBackgroundDetectorLow | 0.2 |
| SLD_lengthTMthr | 5 |
| radiusMedianFilter | 1 |
| minTau | 2 |
| conn3D | 74 |
| estimateOpticalFlow | 0 |
| maxDistPartitionNeigh | 80 |
| deathThrOpticalFlow | -1 |
| minNucleiSize | 8 |
| maxNucleiSize | 3000 |
| maxPercentileTrimSV | 0.2 |
| conn3DsvTrim | 6 |
| maxNumKNNsupervoxel | 10 |
| maxDistKNNsupervoxel | 41 |
| thrSplitScore | -1 |
| thrCellDivisionPlaneDistance | 12.403 |
| thrCellDivisionWithTemporalWindow | 0.456 |

***Table 2: Single guide RNA (sgRNA) sequences for cadherin deletion***

*Pairs of sgRNAs were designed in close proximity to each other to induce large sequence deletions. sgRNAs marked with * are published in (Wu et al., 2018). Italicised sgRNAs were identified as non-functional. The number of base pairs between the sgRNAs Δ was not relevant for cdh2, since their distance was too large for efficient whole-sequence deletion (> 60 kbp).*

| gene | sgRNA 1 | sgRNA 2 | Δ [bp] |
|---|---|---|---|
| cdh1* | GGGATCAAATTACTACCAAC | AGGATTGCTCAAACAGAGGT | 630 |
| cdh11 | GGTGTCTCGAGCCACAGCCT | GCGCATAGTAAGGGCCGTGA | 94 |
| cdh13 | TAACTTGCCACTCAGGTCAG | GGAGCCACGTTACTCTGGGG | 80 |
| cdh15 | AGGGTCATCAAAGTCCACCG | TAGGCTGGAATTTGGGAGCG | 334 |
| cdh17 | ATTGTTCGGGCTGAGGATTT | CTCATTGAGCAGTACGACAC | 38 |
| cdh18a | GCGAGACGTGTTGATTCGTA | CTTGTCGCCGTTGGGCAGTT | 312 |
| cdh2* | *TAAACGATGTACCGTTCCGG* | GGGACTCCAGCCTGGAATGC | --- |
| cdh26.2 | CACTCAGGCGATGGACTACG | ACCGGCACGTTTTTGACGGG | 212 |
| cdh4 | GAATGGGGATGTGAGGCGTC | CGCAGCGGGCATCGACAGAG | 80 |
| cdh5 | TAGTGGCAGAATTCCCAGTC | AATTGGCCTTTTAGAGGTGG | 219 |
| cdh6 | ACGGTGATGTACGTCCTCAC | TTACCAATGTGAGACCCCTC | 235 |
| cdh7a | CGTGCTTCTCGGTCCATGTT | TTGGCCACCACCGACGCCAC | 642 |

***Table 3: Primers used to validate CRISPR gene deletions***

| gene | primer forward | primer reverse |
|---|---|---|
| cdh1 | agtgtcacagggaaatgaaaaca | cctctttcagcaatggttcaga |
| cdh11 | ttcaggggaaggagcaggta | catgaggaacaagctgtcagg |
| cdh13 | tgctgccaattagtctttccg | gcactgagttccatcaaacatca |
| cdh15 | tgtgtttgtgtgcgatcagg | ccaacctccgcaatcagttt |
| cdh17 | tggcatgaacagaaagaagaaca | ctgtggcaagatacatagggt |
| cdh18a | tgtcctctttccttcacacaga | gcccaatgtgctcttccaaa |
| cdh2-1 | cttcttcgctctgacttccg | ccttcaggcaaaagcattcg |
| cdh2-2 | tgatcatccaggctaccgatat | ttaacaacggtgacaaggcc |
| cdh26.2 | cagtgaccttcaaattgcagat | tgagattgaaagaaacaccaggt |
| cdh4 | ggtaacggcctttgacgc | agggtgagtaaacgatgcct |
| cdh5 | gaaaagtgacctcgaccgtg | tgaggcttagcattccatctt |
| cdh6 | cctattgctgcattcccctg | ttcactggtcctcacaagac |
| cdh7a | catacatcattaacggtgcatat | ccttcctgggtatctttgtct |

#### References


Abdelkader, A., Bajaj, C.L., Ebeida, M.S., Mahmoud, A.H., Mitchell, S.A., Owens, J.D., Rushdi, A.A., 2020. VoroCrust. ACM Trans. Graph. 39, 1–16. https://doi.org/10.1145/3337680

Albert, M., 2021. A spatial normalization framework to quantify signaling receptor activity and function during embryonic development. Ruperto Carola University Heidelberg. https://doi.org/10.11588/HEIDOK.00030675

Amat, F., Höckendorf, B., Wan, Y., Lemon, W.C., McDole, K., Keller, P.J., 2015. Efficient processing and analysis of large-scale light-sheet microscopy data. Nat. Protoc. 10, 1679–1696. https://doi.org/10.1038/nprot.2015.111

Ankerst, M., Breunig, M.M., Kriegel, H.P., Sander, J., 1999. OPTICS: Ordering Points to Identify the Clustering Structure. SIGMOD Rec. (ACM Spec. Interes. Gr. Manag. Data). https://doi.org/10.1145/304181.304187

Bukenberger, D.R., Lensch, H.P.A., 2021. Tetrahedra of varying density and their applications. Vis. Comput. 37, 2447–2460. https://doi.org/10.1007/S00371-021-02189-0/FIGURES/17

Ester, M., Kriegel, H.-P., Sander, J., Xu, X., 1996. A Density-Based Algorithm for Discovering Clusters in Large Spatial Databases with Noise, in: Proceedings of the 2nd International Conference on Knowledge Discovery and Data Mining.

Farnsworth, D.R., Saunders, L.M., Miller, A.C., 2020. A single-cell transcriptome atlas for zebrafish development. Dev. Biol. 459, 100–108. https://doi.org/10.1016/J.YDBIO.2019.11.008

Gardiner, J.D., Behnsen, J., Brassey, C.A., 2018. Alpha shapes: determining 3D shape complexity across morphologically diverse structures. BMC Evol. Biol. 18, 184. https://doi.org/10.1186/s12862-018-1305-z

González, Á., 2009. Measurement of areas on a sphere using Fibonacci and latitude-longitude lattices. Math. Geosci. 42, 49–64. https://doi.org/10.1007/s11004-009-9257-x

Hummel, R., 1977. Image enhancement by histogram transformation. Comput. Graph. Image Process. 6, 184–195. https://doi.org/10.1016/S0146-664X(77)80011-7

Jumper, J., Evans, R., Pritzel, A., Green, T., Figurnov, M., Ronneberger, O., Tunyasuvunakool, K., Bates, R., Žídek, A., Potapenko, A., Bridgland, A., Meyer, C., Kohl, S.A.A., Ballard, A.J., Cowie, A., Romera-Paredes, B., Nikolov, S., Jain, R., Adler, J., Back, T., Petersen, S., Reiman, D., Clancy, E., Zielinski, M., Steinegger, M., Pacholska, M., Berghammer, T., Bodenstein, S., Silver, D., Vinyals, O., Senior, A.W., Kavukcuoglu, K., Kohli, P., Hassabis, D., 2021. Highly accurate protein structure prediction with AlphaFold. Nature 596, 583–589. https://doi.org/10.1038/s41586-021-03819-2

Kearse, M., Moir, R., Wilson, A., Stones-Havas, S., Cheung, M., Sturrock, S., Buxton, S., Cooper, A., Markowitz, S., Duran, C., Thierer, T., Ashton, B., Meintjes, P., Drummond, A., 2012. Geneious Basic: An integrated and extendable desktop software platform for the organization and analysis of sequence data. Bioinformatics 28, 1647–1649. https://doi.org/10.1093/bioinformatics/bts199

Kobak, D., Berens, P., 2019. The art of using t-SNE for single-cell transcriptomics. Nat. Commun. https://doi.org/10.1038/s41467-019-13056-x

Lavorini, V., 2018. Gaussian Mixture Model clustering: how to select the number of components. Towar. Data Sci.

Lucy, L.B., 1974. An iterative technique for the rectification of observed distributions. Astron. J. 79, 745. https://doi.org/10.1086/111605

McDole, K., Guignard, L., Amat, F., Berger, A., Malandain, G., Royer, L.A., Turaga, S.C., Branson, K., Keller, P.J., 2018. In Toto Imaging and Reconstruction of Post-Implantation Mouse Development at the Single-Cell Level. Cell 175, 859-876.e33. https://doi.org/10.1016/J.CELL.2018.09.031

McLachlan, G.J., Rathnayake, S., 2014. On the number of components in a Gaussian mixture model. Wiley Interdiscip. Rev. Data Min. Knowl. Discov. 4, 341–355. https://doi.org/10.1002/widm.1135

Nielsen, F., 2019. On the Jensen–Shannon Symmetrization of Distances Relying on Abstract Means. Entropy 21, 485. https://doi.org/10.3390/e21050485

Patel, E., Kushwaha, D.S., 2020. Clustering Cloud Workloads: K-Means vs Gaussian Mixture Model. Procedia Comput. Sci. 171, 158–167. https://doi.org/10.1016/j.procs.2020.04.017

Reynolds, D., 2009. Gaussian Mixture Models, in: Encyclopedia of Biometrics. Springer US, Boston, MA, pp. 659–663. https://doi.org/10.1007/978-0-387-73003-5_196

Richardson, W.H., 1972. Bayesian-Based Iterative Method of Image Restoration. J. Opt. Soc. Am. 62, 55. https://doi.org/10.1364/JOSA.62.000055

Rousseeuw, P.J., 1987. Silhouettes: A graphical aid to the interpretation and validation of cluster analysis. J. Comput. Appl. Math. 20, 53–65. https://doi.org/10.1016/0377-0427(87)90125-7

Sainlot, M., Nivoliers, V., Attali, D., 2017. Restricting Voronoi diagrams to meshes using corner validation. Comput. Graph. Forum 36, 81–91. https://doi.org/10.1111/cgf.13247

Scott, D.W., 1992. Multivariate Density Estimation, 2nd ed, Multivariate Density Estimation: Theory, Practice, and Visualization, Wiley Series in Probability and Statistics. Wiley. https://doi.org/10.1002/9781118575574

Varadi, M., Anyango, S., Deshpande, M., Nair, S., Natassia, C., Yordanova, G., Yuan, D., Stroe, O., Wood, G., Laydon, A., Žídek, A., Green, T., Tunyasuvunakool, K., Petersen, S., Jumper, J., Clancy, E., Green, R., Vora, A., Lutfi, M., Figurnov, M., Cowie, A., Hobbs, N., Kohli, P., Kleywegt, G., Birney, E., Hassabis, D., Velankar, S., 2022. AlphaFold Protein Structure Database: massively expanding the structural coverage of protein-sequence space with high-accuracy models. Nucleic Acids Res. 50, D439–D444. https://doi.org/10.1093/nar/gkab1061

Wang, P., Xin, S., Tu, C., Yan, D., Zhou, Y., Zhang, C., 2020. Robustly computing restricted Voronoi diagrams (RVD) on thin-plate models. Comput. Aided Geom. Des. 79, 101848. https://doi.org/10.1016/j.cagd.2020.101848

Ward, J.H., 1963. Hierarchical Grouping to Optimize an Objective Function. J. Am. Stat. Assoc. 58, 236–244. https://doi.org/10.1080/01621459.1963.10500845

Wit, E., Heuvel, E. van den, Romeijn, J.-W., 2012. 'All models are wrong...': an introduction to model uncertainty. Stat. Neerl. 66, 217–236. https://doi.org/10.1111/j.1467-9574.2012.00530.x

Wu, R.S., Lam, I.I., Clay, H., Duong, D.N., Deo, R.C., Coughlin, S.R., 2018. A Rapid Method for Directed Gene Knockout for Screening in G0 Zebrafish. Dev. Cell 46, 112-125.e4.

https://doi.org/10.1016/j.devcel.2018.06.003

Yan, D.M., Wang, W., Lévy, B., Liu, Y., 2010. Efficient computation of 3D clipped Voronoi diagram. Lect. Notes Comput. Sci. (including Subser. Lect. Notes Artif. Intell. Lect. Notes Bioinformatics) 6130 LNCS, 269–282. https://doi.org/10.1007/978-3-642-13411-1_18/COVER
